# Exhausted-like effector CD8 T cells mediate immune-stromal interactions at mucosal Chronic Graft-versus-Host Disease onset

**DOI:** 10.1101/2025.09.11.675642

**Authors:** Ana Caroline Costa-da-Silva, Rubina Sharma, Marit H. Aure, Daniel Martin, Noemi Kedei, Damilola F. Killanin, Ayesha Javaid, Francis A. Boksa, Drashty P. Mody, Joe T. Nguyen, Sukirth M. Ganesan, Joshua T. Dodge, Clara H. Kim, Marie Kao-Hsieh, David E. Kleiner, Christopher G. Kanakry, Steven Z. Pavletic, Jacqueline W. Mays

**Affiliations:** National Institute of Dental and Craniofacial Research, National Institutes of Health, Bethesda, MD, USA; Collaborative Protein Technology Resource, OSTR, Office of the Director, Center for Cancer Research, National Cancer Institute, National Institutes of Health, Bethesda, MD, USA; Laboratory of Pathology, Center for Cancer Research, National Cancer Institute, National Institutes of Health, Bethesda, MD, USA; Center for Immuno-Oncology, Center for Cancer Research, National Cancer Institute, National Institutes of Health, Bethesda, MD, USA; Immune Deficiency Cellular Therapy Program, National Cancer Institute, National Institutes of Health, Bethesda, MD, USA

## Abstract

Chronic graft-versus-host disease (cGVHD) frequently follows allogeneic hematopoietic stem cell transplantation and impacts the mucosa. This atlas study highlights the role of exhausted-like effector CD8^+^ T cells (T_EXEF_) in oral mucosal cGVHD pathogenesis. Biopsies from the oral mucosa were collected at oral cGVHD onset or at six months post-transplant and analyzed using single-cell RNA sequencing and other modalities. At cGVHD onset, in addition to changes in the myeloid compartment, two distinct populations of CD8^+^T cells with a T_EXEF_ phenotype were prevalent: an inflammatory CD8T cell cluster (CD8T_4) expressing *CCL4*, *CD69*, and *TNFSF9*, and an exhausted CD8T cell cluster (CD8T_5) characterized by the expression of *CXCL13*, *HIF1A*, and *HLA-DRA,* both correlated with clinical disease severity. Pseudo-time analysis suggests a transition from the inflammatory pre-exhausted CD8T_4 cluster to the cytolytic, terminally exhausted CD8T_5 cluster. The CXCL9-CXCR3 interactions among CD8^+^ T cells, fibroblasts and myeloid cells are implicated in tissue damage. Onset of cGVHD despite frequent oral mucosal FOXP3⁺ regulatory CD4⁺T cells suggests potential impaired Treg function or CD8^+^T cell dysfunction. These findings highlight T_EXEF_ cells as key contributors to oral cGVHD pathology and potential therapeutic targets for mitigating sustained tissue injury in cGVHD.

## INTRODUCTION

Chronic graft-versus-host disease (cGVHD) is a significant and multifaceted complication of allogeneic hematopoietic cell transplantation (alloHCT) that impacts mortality and long-term quality of life. Though incidence has decreased, the 2-year cumulative incidence of chronic GVHD requiring systemic treatment remains ∼30% to 40% by National Institutes of Health criteria (1–4). Tissues are heterogeneously impacted; however, the oral cavity is a frequently involved site with a spectrum of oral manifestations including lichenoid lesions, erythema ulceration, mucoceles, immune-mediated salivary gland dysfunction, and progressive tissue sclerosis (5–7). These manifestations not only cause substantial morbidity, including pain, xerostomia, and impaired oral intake, but are also notoriously resistant to conventional immunosuppression. Despite its prevalence and clinical burden, the cellular and molecular drivers of oral cGVHD remain incompletely understood.

In general, the magnitude of the immune response in cGVHD is determined by several factors, including the intensity of initial tissue injury, the proinflammatory chemokine milieu, myeloid cell polarization, and the balance of T-cell subsets. An imbalance between effector and regulatory immune mechanisms has been closely associated with cGVHD development and is a major focus of current research (8, 9). This imbalance, broadly characterized by expansion and differentiation of alloreactive T and B cells into pathogenic subsets, like Th1/Tc1 and Th17/Tc17, and impaired Tregs, sets the stage for chronic inflammation and fibrosis (8–10). Acute inflammation and tissue damage which impair Treg development have been implicated in thymic pathology in cGVHD (8, 9, 11, 12), however, less attention has been paid to how acute inflammation affects other organs, such as the skin, lungs or oral mucosa, despite their frequent involvement in cGVHD. Most insights into these processes derive from preclinical models, underscoring the need for clinical studies to substantiate these findings (13–16).

Different tissues have unique cellular compositions and immune microenvironments that influence their signaling profiles and thereby establish distinct thresholds for disease onset. The buccal oral mucosa (OM), a critical barrier tissue at the entry point of the gastrointestinal tract, is uniquely positioned due to its continual exposure to external antigens – including food particles and microbes - thereby necessitating a specialized cellular environment composed of both immune and non-immune cells. Understanding this distinct role is essential to unravel the specific mechanisms driving cGVHD in the OM, particularly given its role in initiating and sustaining immune responses.

In this study, we sought to elucidate the tissue microenvironment and to identify potential key cell populations involved in OM cGVHD. To this end, we performed single-cell RNA sequencing (scRNA-seq) and multiplex immunohistochemistry (IHC) on OM biopsies from alloHCT recipients with and without clinical evidence of cGVHD and characterized the immune and stromal landscape at single-cell resolution. Our data indicates an imbalance between inflammatory and anti-inflammatory cell populations and reveals dysregulated non-immune cell populations that may contribute to sustained antigen presentation and perpetuation of chronic inflammation. Importantly, our findings provide new insight into the potential role of exhausted-like effector CD8^+^ T cells (T_EXEF_) cells in driving oral cGVHD thereby laying the foundation for future mechanistic studies and therapeutic targeting of the T_EXEF_ cell population.

## RESULTS

### Transcriptional atlas of oral mucosa reflects distinct cell populations and immune activation in cGVHD patients

Based on clinical symptoms, scores and histological findings, patients were divided into 3 groups: no cGVHD (NG), asymptomatic cGVHD (AG), and cGVHD (GV) (Fig. 1). GV patients met NIH diagnostic criteria (17) for oral cGVHD, had a score on the Oral Mucositis Rating Scale, (OMRS) >0 and a positive histopathology score (HistoS) including apoptotic basal keratinocytes and inflammatory infiltration, while AG patients had positive HistoS with no clinical symptoms (Fig. 1A). Consistent with this, both AG and GV patients presented with a significant increase in immune cells detected with IHC (Fig. 1B, C) and flow cytometry (Fig. 1D, E) compared to NG patients (Fig. S1A). Key patient characteristics along with detailed clinical and histopathologic information are provided in Fig. S1B.

**Figure 1:**
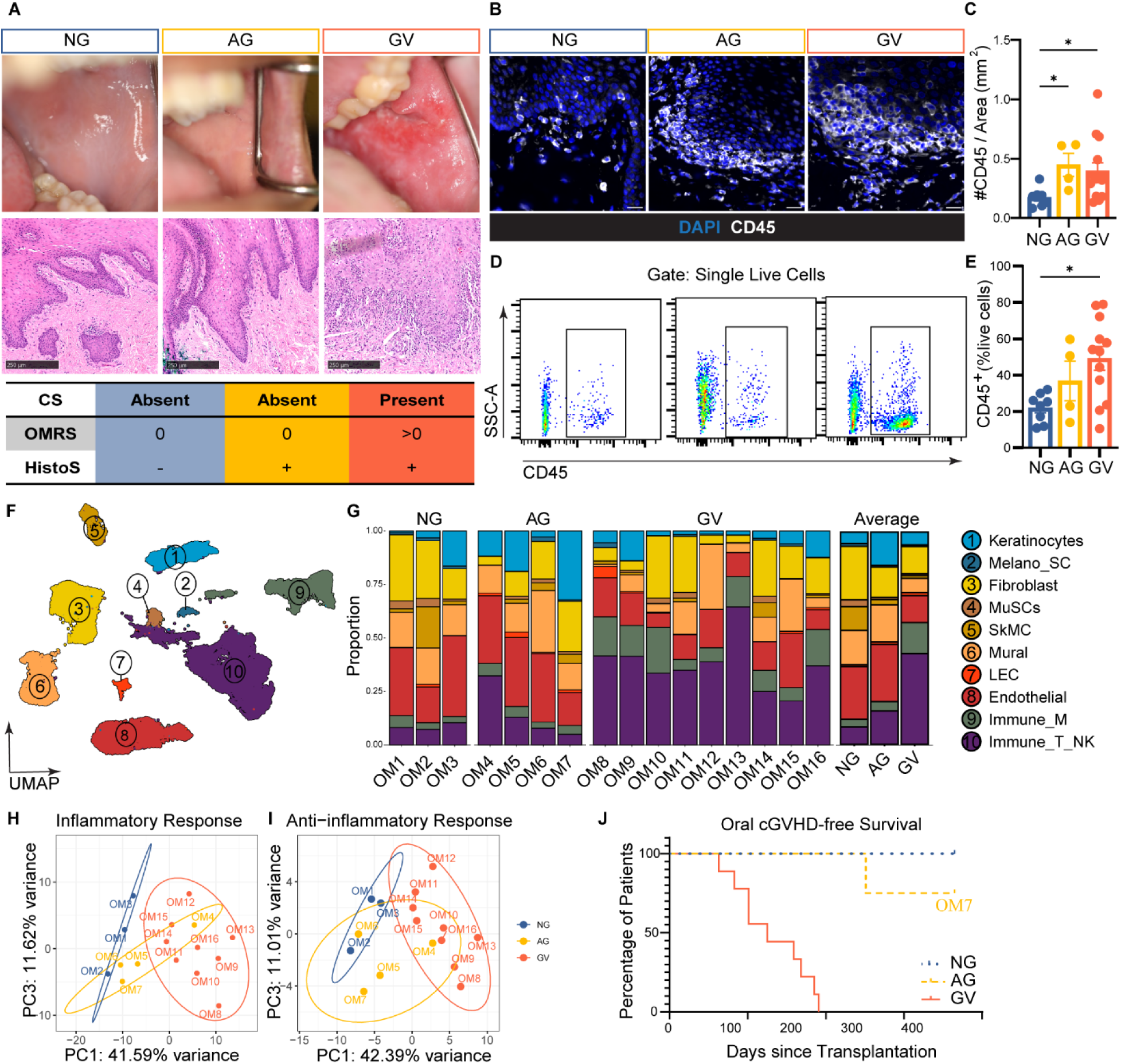
scRNAseq atlas reveals immune activation and heterogeneity in oral cGVHD. **(A)** Representative clinical (top) and histological (middle) images from patients with no oral cGVHD (NG), asymptomatic histopathologic oral cGVHD (AG), and clinically evident oral cGVHD (GV). Clinical severity increases from left to right, with OMRS >0 and HistoS+ indicating GV. H&E staining shows progressive immune infiltration and apoptotic keratinocytes (arrows). Scale bars: 250 µm. **(B)** Immunofluorescence (IF) staining for CD45 (white) in OM of NG, AG and GV patients. DAPI (blue) marks nuclei. Scale bars: 50 µm. **(C)** Quantification of CD45^+^ cells from IF staining for NG (n = 8), AG (n= 4) and GV (n= 14). One-way ANOVA with Kruskal–Wallis correction, ∗p ≤0.01. **(D)** Representative flow cytometry plots showing CD45+ cell gating in NG, AG and GV OM tissue. **(E)** Frequency of CD45⁺ cells among live cells in each group (n=8 NG, n=4 AG, n=12 GV); data are mean ± SEM; one-way ANOVA with Tukey’s correction, *p ≤ 0.01. **(F)** UMAP of 51,617 high-quality OM cells from 16 patients across all groups. Ten transcriptionally distinct clusters were annotated into epithelial (blue), stromal (orange/yellow), and immune (purple/green) compartments. MuSC, Muscle Stem Cell; SkMC, Skeletal Muscle Cells; Mural; LEC, Lymphatic Endothelial Cells; Immune_M, myeloid cells; Immune_T/NK, T and NK cells. **(G)** Stacked bar plots showing the proportion of each cell type by patient (left) and disease group (right). Immune cell proportions are notably expanded in GV. (H-I) Principal Component Analysis (PCA) plots based on inflammatory **(H)** and anti-inflammatory **(I)** gene expression signatures distinguish NG and GV patients, with AG clustering variably. Each point represents one sample. **(J)** Kaplan–Meier analysis of oral cGVHD-free survival curves based on time of oral cGVHD after transplant. x-axis: the time from receiving transplantation to the occurrence of oral cGVHD. y-axis: the probability of oral cGVHD-free survival.

To characterize the cellular composition differences between the patients, we performed scRNAseq on OM from patients from all three groups (NG: n=3, AG: n=4, GV: n=9) (Fig. S1B). After quality control, 51,617 high-quality cells were retained for downstream analysis, comprising 7,932 cells from NG, 8,239 from AG, and 35,446 from GV patients (Fig. S1C). Unsupervised clustering and canonical cell type gene expression profiles revealed 10 distinct OM populations, grouped into three main compartments: epithelium, stroma, and immune (Fig. 1F, Fig. S1D). The epithelial compartment included keratinocytes and melanocytes/Schwann cells (Melano_SC). The stromal compartment was comprised of fibroblasts, muscle stem cells (MuSCs), skeletal muscle cells (SkMC), mural, pericytes, lymphatic endothelial cells (LEC) and endothelial cells. The immune compartment consisted of myeloid cells (Immune_M) and lymphoid cells (Immune/NK) (Fig. 1F).

Proportional comparison of the 10 clusters highlighted heterogeneous distribution among patients in all three groups (Fig. 1G, Fig. S1E). Still, GV had a consistent proportional increase of immune cells, both myeloid (12.52 ± 1.97 % vs 3.83 ± 0.83%) and lymphoid (37.48 ± 4.13% vs 8.63 ± 0.92%;), compared to NG (Fig. 1G). The AG group resembled NG with lower immune cell proportions compared with GV, aligning with their asymptomatic status (Fig. 1A, G). Further, all major cell compartments were identified in the three patient groups, indicating adequate representation in our scRNAseq data.

Principal component analysis (PCA) assessing inflammatory and anti-inflammatory pathway scores showed clear separation of NG and GV groups, while AG had no distinct profile but clustered predominately near NG (Fig. 1H, I). Identification of predictive profiles in AG patients could help define the disease threshold and elucidate cGVHD progression. Notably, one AG patient (OM4) had a mild increased proportion of immune cells and grouped with the GV patients (Fig. 1H, Fig. S1B). However, despite >2 years of clinical follow-up, this patient clustering with GV in the PCA (OM4) never developed clinical signs of oral cGVHD while another AG patient clustering with NG group (OM7) did (Fig. 1H-J). This lack of correlation between inflammatory signatures and onset of oral cGVHD in AG patients highlights the complexity in these cases, which has not been extensively explored in the literature. Thus, further studies in larger cohorts are needed to identify additional factors that may trigger the onset of oral cGVHD, which could be critical for developing predictive models for the disease and refining treatment strategies and diagnostic criteria.

Given the clear separation of disease states between NG and GV samples, we chose to focus our deeper comparative analysis on only these two groups to identify transcriptional hallmarks of disease in non-immune cells, myeloid and lymphoid subpopulations.

### Non-immune cells support the inflammatory niche in oral cGVHD

Non-immune cells are known to play active roles in host defense and pathogenesis by producing immune mediators in various diseases including GVHD (18–20). Non-immune cells in target tissues constitutively express MHC-I and can upregulate MHC-II and co-stimulatory molecules under inflammatory conditions. This upregulation allows them to act as “non-professional” antigen-presenting cells (APCs), initiating, sustaining or amplifying immune responses (21). The role of non-immune OM cells and their transcriptional changes in cGVHD is not well known. Therefore, we sought to identify dysregulated pathways and potential immune mediators in non-immune cells in GV (Fig. 2A). Bulk comparison of non-immune cells identified 2725 differentially expressed genes (DEGs) between NG and GV (Fig. 2B). Among the upregulated genes were inflammatory markers such as *CXCL9, CXCL10, S100A8*, and *HLA-DRA*, while downregulated genes included *ACTA1, DES* and *COX6A*, all related to muscle function (FDR-corrected p-value < 0.05) (Fig. 2B). In line with this, functional analysis revealed significant enrichment in pathways associated with inflammatory responses and chemotaxis in GV, while down-regulated genes were associated with muscle and vascular processes (Fig. 2C). Based on the upregulated pathways, we assigned scores for antigen processing and presentation, chemotaxis and regulation of the inflammatory response. These were particularly prominent in endothelial, fibroblasts, and epithelial clusters suggesting these populations are directly supporting the inflammatory landscape in GV (Fig. 2D-F). To investigate whether specific subpopulations are dysregulated and more involved in GV, we subset and re-clustered the various non-immune groups.

**Figure 2:**
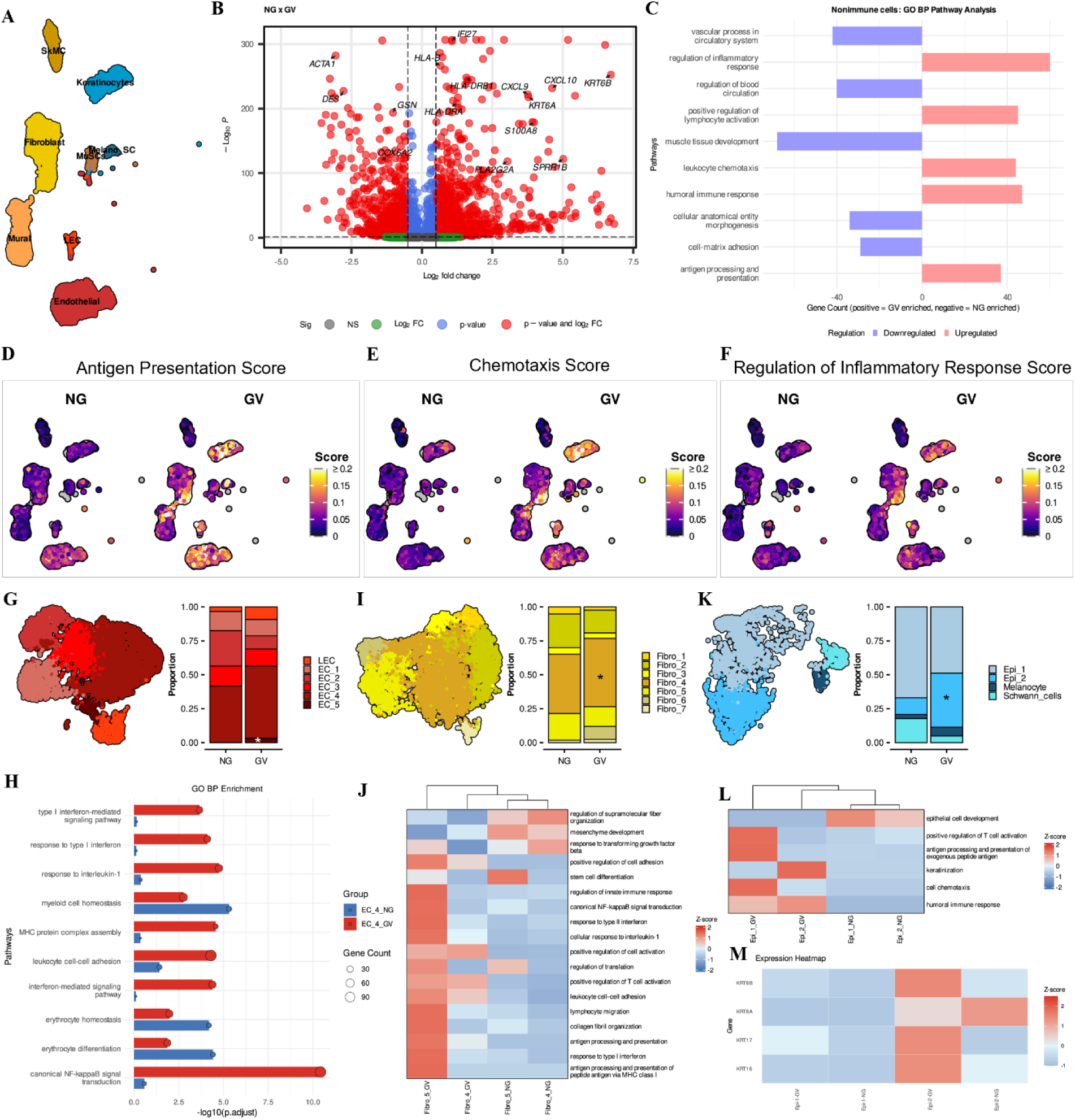
Non-immune cell populations contribute to the inflammatory niche in oral cGVHD. **(A)** UMAP visualization of non-immune cell subclusters identified in NG and GV patients. MuSC, Muscle Stem Cell; SkMC, Skeletal Muscle Cells; Mural; LEC, Lymphatic Endothelial Cells. **(B)** Volcano plot showing DEGs between non-immune cells from GV and NG patients. Analysis performed using MAST; FDR-adjusted *p* < 0.05 and |log₂FC| ≥ 0.5 considered significant. Selected downregulated and upregulated genes in GV patients are highlighted. Nonsignificant genes are represented in blue, green and gray. **(C)** Bar plot of enriched Gene Ontology Biological Process (GO BP) terms in non-immune cells from NG (blue) and GV (pink) patients (FDR < 0.05). **(D-F)** UMAP plots showing UCell enrichment scores for **(D)** antigen presentation, **(E)** chemotaxis, and **(F)** regulation of inflammatory response in non-immune cells from NG and GV. **(G)** UMAP (right) and quantification (left) of endothelial cell clusters in NG and GV samples. Proportional analysis reveals expansion of EC_5 (proliferative) and EC_4 (venous-like) in GV. Two-sided paired Wilcoxon test; p < 0.05. **(H)** Heatmap showing GO BP pathway enrichment on DEGs between EC_4 clusters from NG vs GV patients. Pathways were adjusted using Benjamini–Hochberg method (FDR < 0.05). **(I)** UMAP (right) and cell proportion bar plot (left) of fibroblast clusters correlated to inflammatory process from both NG and GV. Significance was assessed by two-sided paired Wilcox test, and * denotes p < 0.05. **(J)** Heatmap showing GO BP enrichment of DEGs in fibroblast clusters between NG and GV samples. Adjusted p < 0.05 (Benjamini–Hochberg). **(K)** UMAP (right) and proportional analysis (left) of epithelial subclusters from NG and GV patients. Epi_2 (suprabasal keratinocytes) was significantly expanded in GV. **(L)** GO BP enrichment heatmap comparing DEGs in keratinocyte subpopulations (Epi_1 and Epi_2) between NG and GV patients. Pathways linked to chemotaxis and immune activation were significantly enriched in GV. Pathways were adjusted using Benjamini–Hochberg method (FDR < 0.05). **(M)** Scaled expression heatmap of selected genes (e.g., KRT6B, KRT16, KRT17) upregulated in keratinocytes from GV patients, associated with epithelial damage and regeneration.

Muscle tissue is often coincidentally captured at the base of the buccal mucosal punch biopsy and was identified in this dataset. Downregulation of muscle related genes and related pathways suggested detrimental changes to muscle cell types in GV. Sub setting and re-clustering of muscle cells categorized two subtypes of SkMCs, type1-2 (Fig. S2A). There was a high proportional variation of MuSC and type1 SkMCs within the GV group, while type 2 SkMCs were significantly decreased in GV compared to NG (Fig. S2C). Pathway analysis identified several metabolic processes as specifically to type2 SkMCs, suggesting a dysregulation of these in GV (Fig. S2C). However, further work is needed to understand the significance of dysregulated muscle populations in OM GV. Similarly, mural cells were subseted and re-clustered resulting in four distinct populations, smooth muscle cells (SMC) SMC_1-3 and pericytes (Fig. S2D). There were no significant proportional differences of these clusters in GV compared to NG (Fig. S2E). Still, pathway enrichment analysis highlighted the possible contribution of SMC_3 in ECM-related processes as well as chemotaxis and response to type 1 interferon (Fig. S2F). Further comparisons between NG and GV showed that SMC_3 was enriched for genes linked to antigen presentation and response to interferon-gamma (IFNγ), suggesting involvement of this subpopulation in GV (Fig. S2G).

Among endothelial populations we identified five clusters (EC): arterial_EC (EC_1), capillary_EC (EC_2), Tip-like_EC (EC_3), venous_EC (EC_4), and proliferative_EC (EC_5) in addition to one LEC population (Fig. 2G, Fig. S3A). Proportionally, EC_5 was significantly increased in GV patients, suggesting angiogenesis or tissue repair in GV (Fig. 2G, Fig. S3B). Functional characterization revealed enrichment of pathways involved in antigen processing and presentation and regulation of inflammatory responses in the EC_4 cluster and this was upregulated in GV (Fig. S3C). These findings suggest active remodeling of vascular niches to support immune infiltration and tissue damage. These findings were positively correlated with the increased infiltration of immune cells in GV patient tissues.

We identified seven distinct fibroblast subpopulations Fibro_1-7 (Fig. 2I, Fig. S3D). Proportionally, only Fibro_4 cluster was significantly expanded in GV (Fig. 2I, Fig. S3E). Go enrichment analysis showed that Fibro_4 was specifically enriched for cytoplasmic translation, suggesting a specific role in synthesizing proteins while both Fibro_5 and Fibro_6 clusters were more strongly associated with inflammatory pathways, particularly in GV patients (Fig. 2J, Fig. S3F). Notably, Fibro_6 cluster was specifically enriched in one GV patient with the highest OMRS score and further work is needed to understand the role of this subpopulation in GV (Fig. S3E). Overall, these results show that specific subpopulations of fibroblasts displayed a stronger association with inflammatory pathways, and these were elevated in GV compared to NG, highlighting active roles in the heightened inflammatory landscape. Further work is needed to understand whether these findings are tissue specific or if fibroblast subpopulations have a broader role in cGVHD.

Severe oral cGVHD is known to increase epithelial apoptosis in intraepithelial infiltrates (22). However, transcriptional changes associated with cGVHD in OM epithelia are not known. In our dataset, the epithelial compartment consisted of four distinct populations, basal keratinocytes (Epi_1), suprabasal keratinocytes (Epi_2), melanocytes, and schwann cells (Fig. 2K, Fig. S3G). While Epi_1 showed no significant proportional difference between NG and GV (Fig. 2K, Fig. S3H), pathway analysis uncovered a distinct enrichment of genes associated with chemotaxis in GV (Fig. 2L). In contrast, the Epi_2 cluster was significantly enriched in GV patients (Fig. 2K, Fig. S3H), displaying a global gene expression profile linked to leukocyte chemotaxis and antimicrobial immune responses (Fig. 2L and Fig. S3I). Additionally, Epi_2 was enriched for *KRT6B, KRT16*, and *KRT17* and this was exacerbated in GV compared to NG (Fig. 2M). These keratins are known to be damage-associated and play a central role in the process of epithelial damage regeneration (23–26), pointing to a profile of epithelial barrier disruption in GV OM.

Taken together, our data illustrates that subpopulations of fibroblasts, endothelial and epithelial cells, are active participants in sustaining chronic antigen presentation and inflammation and thereby support the inflammatory niche in oral cGVHD.

### Myeloid clusters shape the inflammatory landscape in oral cGVHD

Myeloid cells are critical regulators of the inflammatory landscape and play a pivotal role in initiating and modulating immune responses during GVHD. We bioinformatically isolated and re-clustered myeloid cells and based on canonical markers, we identified mast cells, plasmacytoid dendritic cells (pDCs), four subsets of conventional dendritic cells (cDC_1-4), monocytes (Mono) and four subsets of macrophages (Mac_1-4; Fig. 3A, B and Fig. S4A).

**Figure 3:**
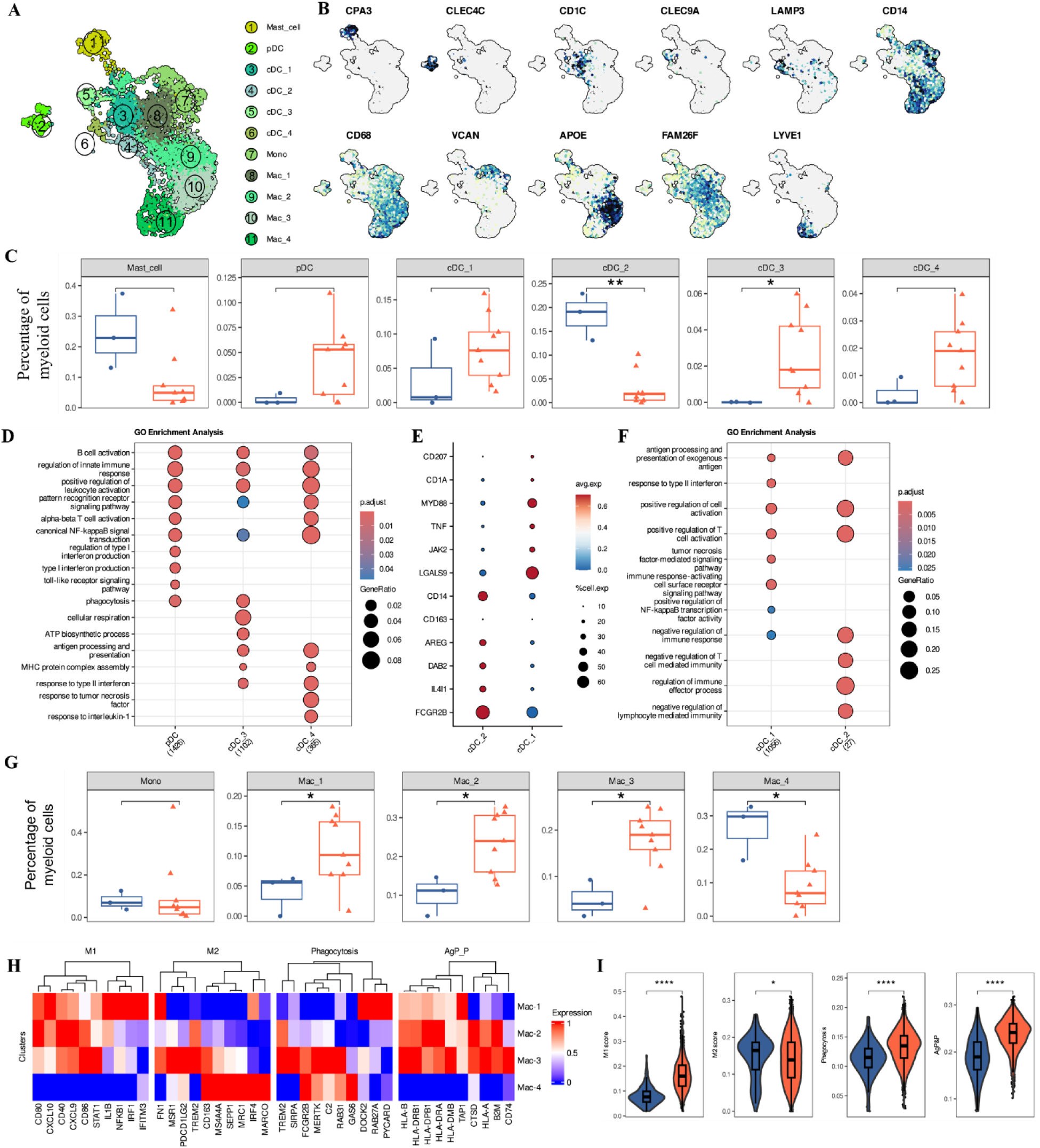
Myeloid cell compartment remodeling sustains an inflammatory microenvironment in oral cGVHD. **(A)** UMAP projection of 5,685 integrated myeloid cells from NG and GV oral mucosa, grouped into 11 transcriptionally distinct clusters. **(B)** Feature plots showing normalized expression of key marker genes used to define dendritic cell (DC), mast cell, monocyte, and macrophage subsets. **(C)** Boxplots depicting the relative abundance of mast cells and dendritic cell subsets in NG (blue) and GV (red) patients. Notably, pDCs, CLEC9A⁺ cDC_3, and LAMP3⁺ cDC_4 subsets are enriched in GV. Statistical significance determined by two-sided Wilcoxon rank-sum test. **(D)** Gene Ontology (GO) Biological Process enrichment of genes upregulated in the three GV-associated DC clusters (pDCs, cDC_3, and cDC_4). Circle size indicates gene ratio; color represents adjusted *p*-value (Benjamini–Hochberg correction; one-sided Fisher’s exact test).**e,** Dot plot showing the mean expression of genes that differentiate cDC_1 and cDC_2. The dot size indicates the percent of expressing cells, and the dot color is the scaled average expression. **(F)** GO BP enrichment of upregulated genes in cDC_1 and cDC_2 clusters, highlighting functional differences between conventional DC subsets. The size of each circle represents the gene ratio, and the color indicates p-adjust using one-sided Fisher’s exact test with BH multiple-testing correction. **(G)** Boxplots showing the relative frequencies of monocyte and macrophage subpopulations in NG and GV patients. Inflammatory macrophage subsets are expanded in GV. Statistical analysis by two-sided Wilcoxon rank-sum test. **(H)** Heatmap showing scaled expression of canonical genes associated with M1, M2, phagocytosis, and antigen processing and presentation (AgP_P) programs across macrophage clusters. **(I)** Violin plots display calculated signature AUCell scores for M1, M2, phagocytosis, and AgP&P pathways in macrophages from NG and GV samples. Signature differences indicate a skewing toward pro-inflammatory and antigen-presenting phenotypes in GV. Statistical comparisons by Wilcoxon rank-sum test.

Proportional heterogeneity was observed within the GV patients, especially among cDC clusters (Fig. S4B). Still, inflammatory-related clusters indicative of DC activation, such as pDCs, CLEC9A^+^XCR1^+^ cDC_3, and LAMP3^+^CCR7^+^ cDC_4, were enriched in GV patients (Fig. 3C, Fig. S4B). Pathway analysis showed these clusters were associated with IFN type 1 signaling pathway, lymphocyte activation and antigen processing and presentation, indicating that they may functionally contribute to the heightened inflammatory response observed in these patients. (Fig. 3D). We also noted a trend of increased proportion of cDC_1 and significant decrease of cDC_2 in GV versus NG patients (Fig. 3C). Both clusters express *CD1C, CLEC10A* and *FCER1A* (Fig. S 4C), but cDC_1 is distinguished by enrichment of markers associated with DC activation signals in tissue (27), while cDC_2 displayed elevated levels of *CD14* and regulatory molecules such as *DAB2* and *AREG*, potentially indicative of a tolerogenic phenotype (27) (Fig. 3E). This was supported by enrichment of pathways related to the tumor necrosis factor (TNF)-mediated signaling and response to type II interferons (IFN-γ) in cDC_1 while cDC_2 was associated with pathways involved in the negative regulation of the immune response, highlighting its potential role in immune modulation (Fig. 3F). These results underscore a shift in dendritic cell composition and function in GV patients, with cDC_1 potentially amplifying immune activation and cDC_2 assuming a regulatory, tolerogenic role—highlighting their possible contribution to disease pathogenesis.

Among macrophage clusters, Mac_1-3 were significantly elevated while Mac_4 was significantly reduced and monocytes were unchanged in GV patients, suggesting a broader shift in immune regulation (Fig. 3G, Fig. S4B). This was further reflected by significantly increased M1-, phagocytosis and antigen processing and presentation gene set scores for Mac_1-2 (Fig. 3H, I, Fig. S4D), reflecting the overall putative functional differences between NG and GV with increased inflammatory polarization (Fig. 3H, I). The Mac_2-3 clusters had DEGs related to lipid metabolism and phagocytosis (*APOE, APOC1;* Fig. S4A). Notably, compared to Mac_2, Mac_3 macrophages also demonstrated higher expression of genes involved in iron metabolism and phagocytosis of apoptotic cells (Fig. S4E). Conversely, Mac_4 had unique expression of *LYVE1* (Fig. 3B) suggesting a tissue-resident subtype along with MRC1 showing highest M2–score, reduced phagocytosis and antigen-processing and presentation scores (Fig. 3H, I, Fig. S4D), consistent with a more reparative role for this cluster. The Mac_3 cluster displayed a mixed profile, combining both M1 and M2 signatures (Fig. 3H), elevated phagocytosis and antigen-processing and presentation scores, suggesting an intermediate macrophage population. When comparing these functional scores across macrophage populations between NG and GV patients, GV patients exhibited higher M1-, phagocytosis-, and antigen-processing and presentation-related scores, and a lower M2 score (Fig. 3I). In summary, these findings highlight how shifts in the composition and functional potential of the myeloid milieu contribute to immune dysregulation and may directly influence the disease outcome.

### Expanded T_EXEF_ Cells Accumulate at the Epithelial Basement Membrane Alongside Tregs in cGVHD

T cells exhibit significant heterogeneity in cellular composition and functional states in cGVHD and are critical for understanding tissue specific damage. Flow cytometric analysis showed that, compared to NG, GV patients presented a significant overall increase in total CD8⁺T cells, while CD4⁺ T cell frequencies remained comparable (Fig. S5A). To explore transcriptional changes within specific subpopulations, T and NK cells were bioinformatically re-clustered and, based on known markers, eight distinct subpopulations were identified; CD4T_1 (*IL17R^+^, ANXA1^+^),* CD4T_2 (regulatory T cell, Treg: *FOXP3^+^, IL2RA^+^*), NK_gdT (natural killer– unconventional T cell, expressing markers characteristic of both NK cells and unconventional T cells: *GNLY^+^,TRDC^+^*), CD8T_1 (proliferative, *TOP2A^+^, MKI67^+^*), CD8T_2 (effector memory *GZMK*^high^), CD8T_3 (*IL7R^+^*), CD8T_4 and CD8T_5 (exhaustion, *LAG3^+^, HAVCR2^+^*) (Fig. 4A, B and Fig. S5B).

**Figure 4:**
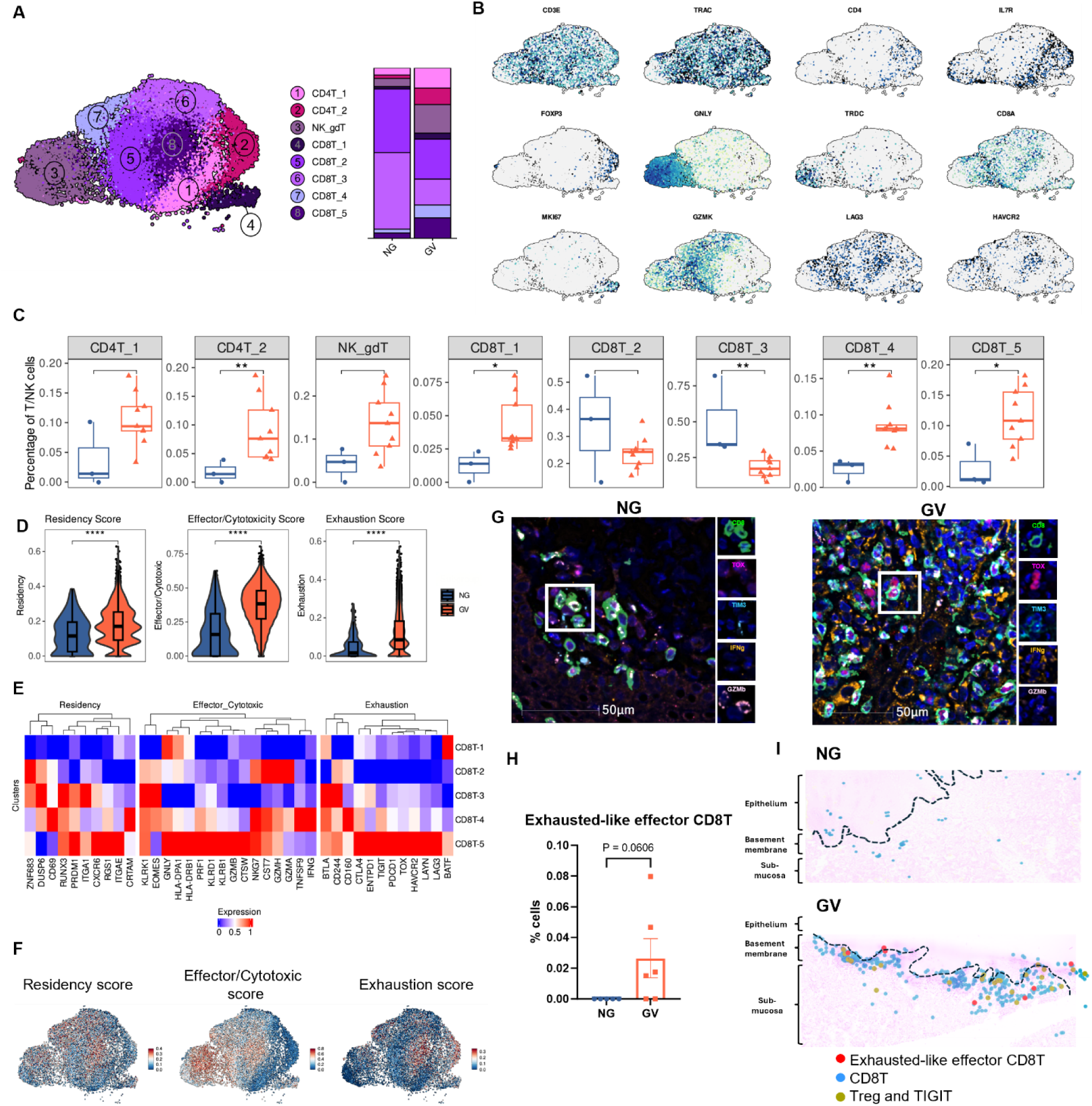
Mapping of the T_EXEF_ cell phenotype. **(A)** (right) UMAP projection of 16,692 lymphoid cells from NG and GV oral mucosa samples, grouped into eight transcriptionally distinct clusters; (left) Stacked bar plots showing the proportion of each cell type by disease group (right). **(B)** Feature plots showing normalized expression of representative genes used to define CD4⁺ T cells, CD8⁺T cells, NK cells, and regulatory T cell subsets. **(C)** Boxplots showing the relative proportions of lymphoid cell subsets in NG (blue) and GV (red) patients. Statistical comparisons performed using the Wilcoxon rank-sum test. **(D)** Violin plots showing AUCell signature scores (residency, effector/cytotoxic, and exhaustion) across combined CD8T_2, CD8T_3, and CD8T_4 clusters in NG and GV patients. Significant enrichment of exhaustion-associated signatures observed in GV. *p*-values calculated via Wilcoxon rank-sum test. **(E)** Heatmap showing scaled signature scores for tissue residency, cytotoxicity, and exhaustion across CD8⁺T cell clusters. **(F)** UMAP visualization of the spatial distribution of same AUCell signature scores shown in (E), highlighting heterogeneity across CD8⁺ T cell populations. **(G)** Multiplex immunofluorescence staining of OM tissue showing co-localization of CD8 (green), TOX (magenta), TIM-3 (cyan), IFNγ (orange), and GZMB (light pink); DAPI (blue) marks nuclei. Images acquired at 20× magnification; scale bars = 50 µm. **(H)** Quantification of CD8⁺TOX^+^Tim-3^+^IFNγ^+^GZMB^+^ T cells shown in **(G)** in NG (n = 5) and GV (n = 6) samples. *p*-values determined using Wilcoxon rank-sum test. **(H)** Spatial mapping overlay of representative NG (top) and GV (bottom) OM tissues showing T_EXEF_ (red), conventional CD8⁺ T cells (blue), and TIGIT⁺ regulatory T cells (olive-green), overlaid on H&E-stained sections. Black line indicates epithelial boundary.

There was a significant increase of FOXP3⁺ CD4⁺Tregs (CD4T_2), proliferative CD8⁺ T cells (CD8T_1), and the exhausted subsets CD8T_4 and CD8T_5. In contrast, CD8T_3 cells were significantly decreased in GV (Fig. 4A, C and Fig. S5C). Notably, CD8T_4 and CD8T_5 were almost exclusively found in GV samples (Fig. 4A, C and Fig. S5C). To better define the functional state of CD8⁺T cells, we compared enrichment scores for tissue residency, cytotoxicity, and exhaustion using established gene sets (see Materials & Methods). All three scores were significantly elevated in GV samples (Fig. 4D). Similarly, we assessed and compared the same enrichment scores across CD8 subpopulations. CD8T_4 and CD8T_5 exhibited significantly higher scores across all categories, indicating a highly activated, resident, and exhausted phenotype (Fig. 4E and Fig. S5D). CD8T_2 resembled tissue-resident effector memory cells, while CD8T_3 expressed quiescent/resting-associated genes (Fig. 4E, f; Fig. S5D).

Validation by multiplex immunostaining confirmed the presence of CD8⁺T cells co-expressing TIM3, TOX, IFNγ, and GZMB — hallmark features of the exhausted-like effector phenotype — primarily in GV samples (p = 0.06; Fig. 4G, H). A significant increase in the frequencies of CD8⁺ T cells individually expressing these markers were also observed in GV (Fig. S5E, F). Supporting the scRNA-seq findings, multiplex IHC also showed a significant enrichment of TIGIT⁺ CD4⁺Tregs (CD4T_2) in GV oral mucosa compared to controls (p = 0.02; Fig. S5G). Spatial analysis revealed that these TIGIT⁺ Tregs were often located near CD8⁺ T cells, suggesting possible regulatory interactions within the GV tissue microenvironment (Fig. 4I). Importantly, CD8⁺ T cells including, T_EXEF_ cells, were localized near the epithelial layer and adjacent to the basement membrane — key sites of immune infiltration and tissue damage in GV pathology (Fig. 4I). Taken together, these findings highlight a coordinated accumulation of T_EXEF_ cells and regulatory CD4⁺T cells at key epithelial sites in OM during cGVHD, suggesting that while Tregs are enriched, their presence appears to be insufficient to suppress pathogenic CD8⁺T cell activity and limit tissue damage.

### Functional Specialization and Developmental Trajectory of T_EXEF_ Cell Subsets in cGVHD

To better understand how T_EXEF_ cells contribute to cGVHD pathogenesis, we analyzed DEGs of the CD8T_4 and CD8T_5 clusters, both almost exclusively present in GV patients. Compared to all CD8+ T cells, this analysis revealed 920 upregulated genes (Fig. 5A), providing insight into the unique transcriptional programs associated with these two T_EXEF_. CD8T_4 and CD8T_5 shared 56 upregulated genes (6.1%), including *IFNG, GZMB,* and *HAVCR2* (Fig. 5A, B). Interestingly, *IFNG* expression was higher in CD8T_4, suggesting a more pro-inflammatory effector state (Fig. 5B). In contrast, *GZMB* and *HAVCR2* were highly expressed in CD8T_5, indicating a more cytotoxic and terminally exhausted-like phenotype (Fig. 5B). GO biological process analysis of upregulated genes identified that 2327 pathways (46.1%) were shared between the two clusters (Fig. 5C), including pathways involved in cytotoxicity and regulation of T cell activation (Fig. 5D). CD8T_4-specific pathways included responses to TNF and innate immune regulation, that are consistent with heightened inflammatory reactivity or stress-responsiveness (Fig. 5D). CD8T_5-specific pathways pointed to processes including MHC class II complex assembly, post-translational protein modification, and ATP synthesis (Fig. 5D), that suggest metabolic and transcriptional reprogramming. These data support a model of functional specialization between the two subsets: CD8T_4 appears to represent a more inflammatory, pre-exhausted population, while CD8T_5 reflects a metabolically adapted, terminally exhausted-like state. This was further supported by cluster-specific gene expression: *CCL4, CD69,* and *TNFSF9* were enriched in CD8T_4, while *CXCL13, HIF1A,* and *HLA-DRA* were upregulated in CD8T_5 (Fig. 5B). The clinical OMRS score positively correlated with the expression of CCL4 in the CD8T_4 cluster (Fig. 5E) and CXCL13 in the CD8T_5 cluster (Fig. 5F) in this cohort, indicating that these chemokine-defined exhausted-like subsets are not only transcriptionally distinct but may indicate clinical severity and disease activity in oral cGVHD.

**Figure 5.**
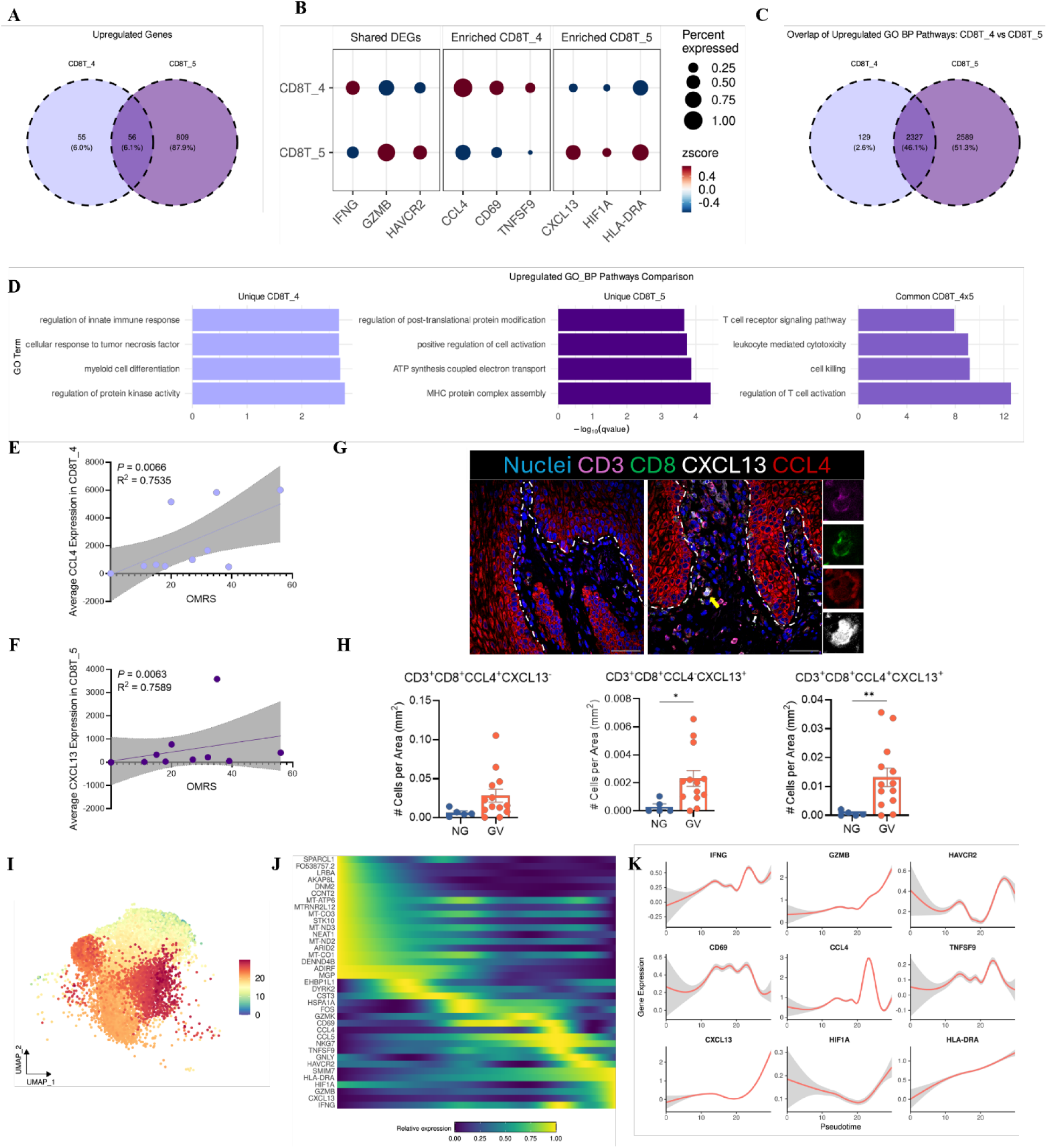
Shared and distinct transcriptional programs define terminally differentiated CD8⁺ T cell subsets in oral cGVHD. **(A)** Venn diagrams showing differentially expressed genes (DEGs) upregulated in CD8T_4 and CD8T_5 clusters compared to other CD8⁺ T cell subsets. Left: shared DEGs; middle: CD8T_4-specific; right: CD8T_5-specific. DEGs identified using MAST with FDR-adjusted *p* < 0.05 and |log₂FC| ≥ 0.5. **(B)** Dot plot displaying expression of upregulated genes shared by or enriched in CD8T_4 and CD8T_5 clusters. Dot size represents the percentage of cells expressing gene; color indicates average expression. **(C)** Venn diagrams showing the numbers of overlapping and non-overlapping pathways for upregulated DEGs shared (left), unique to CD8T_4 (middle) and CD8T_5 (right) between CD8T_4 and CD8T_5 clusters. Pathways shown are significant at FDR < 0.05. **(D)** Barplots show selected GO BP pathway pathways for upregulated DEGs unique and shared between CD8T_4 and CD8T_5 clusters. All pathways shown were significantly enriched at FDR < 0.05. Correlation plots comparing CCL4 gene expression in CD8T_4 cluster **(E)** and CXCL13 gene expression in CD8T_5 cluster **(F)** with the Oral Mucositis Rating Scale (OMRS). Each data point represents one patient. The solid line represents the best fit and the gray region represents its 95% confidence intervals. **(G)** Representative multiplex immunofluorescence staining of oral mucosa tissue showing CD3 (magenta), CD8 (green), CXCL13 (white), and CCL4 (red). DAPI (blue) labels nuclei. Images captured at 20× magnification; scale bars = 50 μm. **(H)** Quantification of CD8⁺ T cells co-expressing CCL4 alone, CXCL13 alone, or both CCL4 and CXCL13 in NG (n = 5) and GV (n = 13) oral mucosa samples. Bars indicate mean ± SEM. Statistical comparisons performed using the Wilcoxon rank-sum test; *p* < 0.05 considered significant. **(I)** Diffusion pseudotime analysis using Slingshot across CD8⁺ T cell clusters (excluding proliferating cells). Pseudotime is color-coded from early (blue) to late (red), indicating a transition from naïve to terminally exhausted states. **(J)** Heatmap showing dynamic expression profiles of pseudotime-associated genes (adjusted *p* < 0.01, tradeSeq association test), ordered by increasing pseudotime. Rows represent genes; columns represent single cells color-coded by cluster identity. **(K)** Expression trajectories of selected exhaustion- and effector-associated genes as a function of pseudotime, visualized across the inferred CD8⁺ T cell differentiation continuum.

The differential expression of CCL4 and CXCL13 prompted investigation of their protein-level expression via immunohistochemistry. We observed an increasing trend of CCL4⁺CXCL13⁻ CD8⁺ T cells in GV samples, alongside significant increases in frequency of both CCL4⁺CXCL13⁺ and CCL4⁻CXCL13⁺ subsets (Fig. 5G, H), suggesting potential transitional states between pre-exhausted and terminally exhausted phenotypes. To better understand the dynamic changes in T cells in cGVHD, we applied pseudotime trajectory analysis on non-proliferative CD8 T cells and visualized it in UMAP space (Fig. 5I). The heatmap shows dynamic expression changes of DEGs, within CD8 T cells (Fig. 5J), and distribution patterns of named genes confirmed the sequential upregulation of key markers (Fig. 5K). *CCL4, CD69, TNFSF9,* and *IFNG* appeared earlier, aligning with CD8T_4, while *CXCL13, HAVCR2, GNLY,* and *GZMB* increased later, defining the CD8T_5 subset (Fig. 5J, K). Interestingly, *IFNG* displayed a fluctuating pattern, potentially reflecting dynamic transitions in effector function during exhaustion. In summary, CD8T_4 and CD8T_5 represent distinct T_EXEF_ cell states in cGVHD, characterized by specific transcriptional, functional, and developmental profiles. Their spatial and temporal dynamics suggest a continuum from inflammatory pre-exhaustion to terminal cytolytic exhaustion, with implications for targeted immune modulation.

### Altered cell crosstalk in cGVHD involves T_EXEF_ cells

Cell interaction analysis highlighted significant alterations in general immune crosstalk patterns that had an increased number of interactions and interaction strength in GV (Fig. S6A). Specifically, the CD8T_4 T_EXEF_ subpopulation had increased importance in incoming signaling in GV, while in NG, the incoming signal of importance was through the anti-inflammatory Mac_4 (Fig. S6B) pointing to a shift in cell interactions towards damage in GV.

We then employed Tensor-cell2cell on LIANA-inferred ligand-receptor scores to further investigate context-specific intercellular communication differences between NG and GV samples, focusing on the two populations of interest – CD8T_4 and CD8T_5 (18). We first analyzed their interactions with non-immune cells and observed communication contexts enriched in both GV (Fig. 6A) and NG (Fig. S6A) patients. Enriched context in GV involved fibroblasts, pericytes, and SMC_3 (Fig. 6B) and it was mainly controlled by upregulation of the TGFβ signaling pathway, as shown by PROGENy-based activity scores (Fig. 6C). Within the TGFβ pathway, there was a pronounced increase in fibroblast-CD8⁺ T cell communication, particularly through extracellular matrix-integrins interactions including *FN1* and *ITGA4–TGB1* known to be involved in cell migration (Fig. 6D). When analyzing significantly decreased interactions in GV samples related to CD8T cells and non-immune populations (Fig. S6C), we observed frequent participation of epithelial, fibroblasts and endothelial cells (Fig. S6D). Downstream pathway analysis also showed TGFβ as the top activated pathway (Fig. S6E) Interestingly, we identified high contribution of extracellular matrix (ECM)-receptor signaling interactions, such as *COL1A1– SDC1*, *THBS1–SDC1*, and *TNC–CD44* withing TGFβ pathway (Fig. S6F).

**Figure 6:**
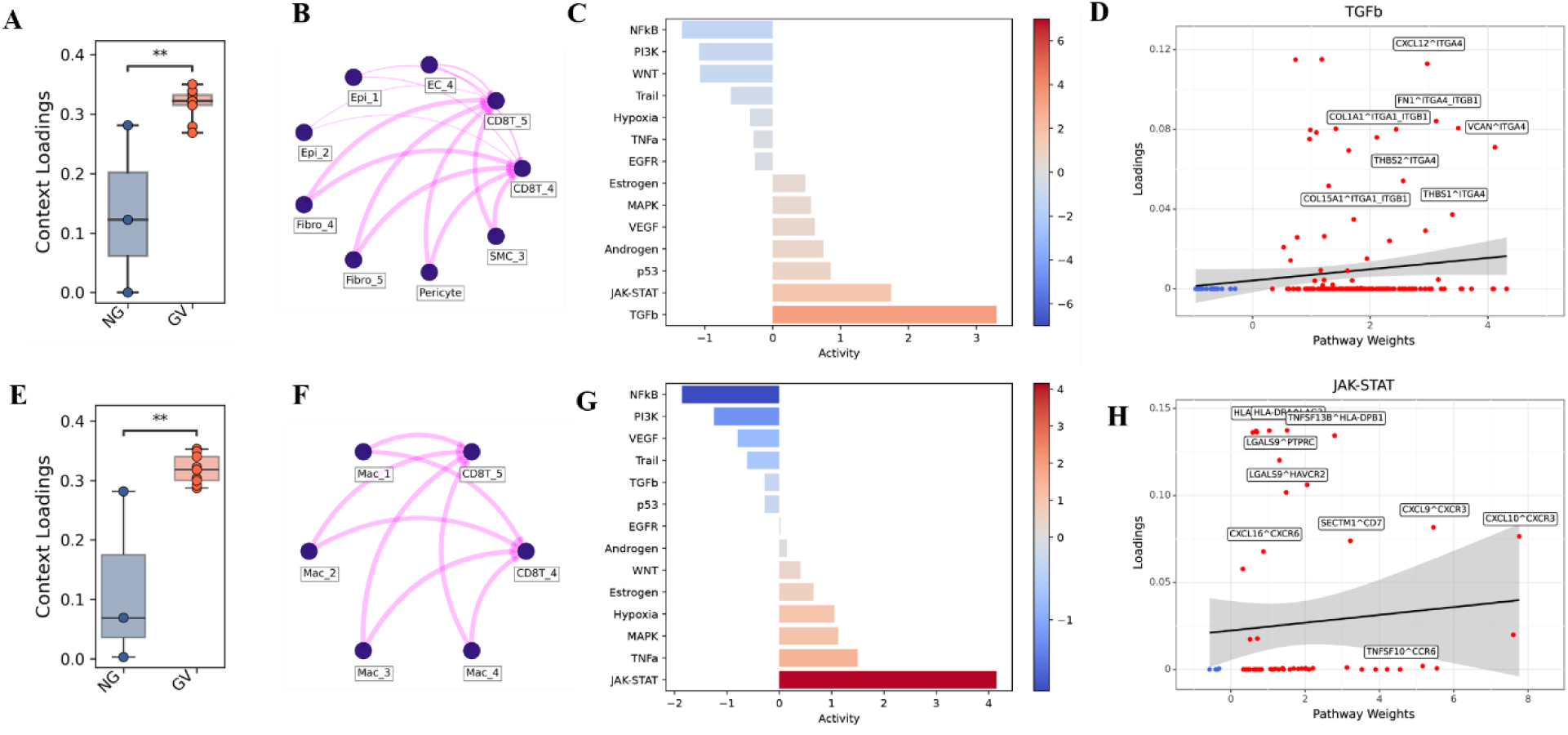
Cell–cell communication and signaling network alterations in oral mucosa during chronic GVHD. **(A)** Box plots compare upregulated context loadings between NG and GV patients related to stromal/epithelial-CD8^+^ T cells interactions identified through Tensor-cell2cell analysis. Pairwise t-tests were performed, and p-values were adjusted using the Benjamini– Hochberg method. Statistical significance is indicated by asterisks (p < 0.01). **(B)** Network diagrams illustrate interactions between CD8⁺ T cells and stromal/epithelial cells corresponding to (A), with connecting lines representing interaction strength. **(C)** Pathway enrichment analysis of ligand–receptor loadings identified using PROGENy. Color intensity reflects activity magnitude and reveals elevated TGF-β signaling activity in GV patients. **(D)** Scatter plots showing ligand– receptor pairs contributing to the TGF-β pathway enrichment shown in (C), with regression lines and confidence intervals (shaded areas) indicating trends and variability. **(E)** Box plots compare upregulated context loadings between NG and GV patients related to macrophages-CD8^+^ T cells interactions identified through Tensor-cell2cell analysis. Pairwise t-tests were performed, and p-values were adjusted using the Benjamini–Hochberg method. Statistical significance is indicated by asterisks (p < 0.01). **(F)** Network diagrams illustrate interactions between CD8⁺ T cells and macrophages corresponding to (E), with connecting lines representing interaction strength. **(G)** Pathway enrichment analysis of ligand–receptor loadings identified using PROGENy. Color intensity reflects activity magnitude and reveals elevated JAK-STAT signaling activity in GV patients. **(H)** Scatter plots showing ligand–receptor pairs contributing to the JAK-STAT pathway enrichment shown in (G), with regression lines and confidence intervals (shaded areas) indicating trends and variability.

Regarding interactions between CD8T clusters and myeloid cells, we observed significant enrichment context in GV involving both macrophages (Fig 6E, F) and dendritic cells (Fig. S6G, H). In both cases, JAK-STAT signaling was the top activated pathway (Fig. 6G and Fig. S6I), primarily involving chemokine signaling including the known interaction between *CXCL9– CXCL10* to *CXCR3* (Fig. 6H and Fig. S6J). Unique to macrophage - CD8T cell crosstalk was a prominent *CXCL16–CXCR6* signaling axis, that originated from macrophages and included interactions with both CD8T_4 cells and CD4T_2 populations (Fig. S6K).

Our integrated single-cell and spatial profiling of oral mucosa in cGVHD reveals a coordinated network of immune-stromal interactions that sustains chronic CD8⁺ T cell activation and tissue damage. We propose a working model where epithelial barrier disruption and persistent antigen presentation drive the emergence of two functionally distinct T_EXEF_ cell subsets: an inflammatory pre-exhausted population (CD8T_4) and a terminal cytotoxic subset (CD8T_5). Stromal cells, including fibroblasts, epithelial, and endothelial cells, actively contribute to the inflammatory niche by expressing antigen-presenting and chemotactic molecules. Concurrently, myeloid cells (e.g., LAMP3⁺ dendritic cells and APOE⁺ macrophages) engage CD8⁺ T cells through CXCL16–CXCR6 and IFN-mediated signaling. This cellular crosstalk facilitates spatial accumulation of effector T cells at epithelial interfaces, where tissue destruction persists despite increased presence of Tregs (Fig. 7), These findings provide a mechanistic framework for how chronic immune activation is sustained in oral cGVHD and identify potential cellular targets for future therapeutic intervention.

**Figure 7:**
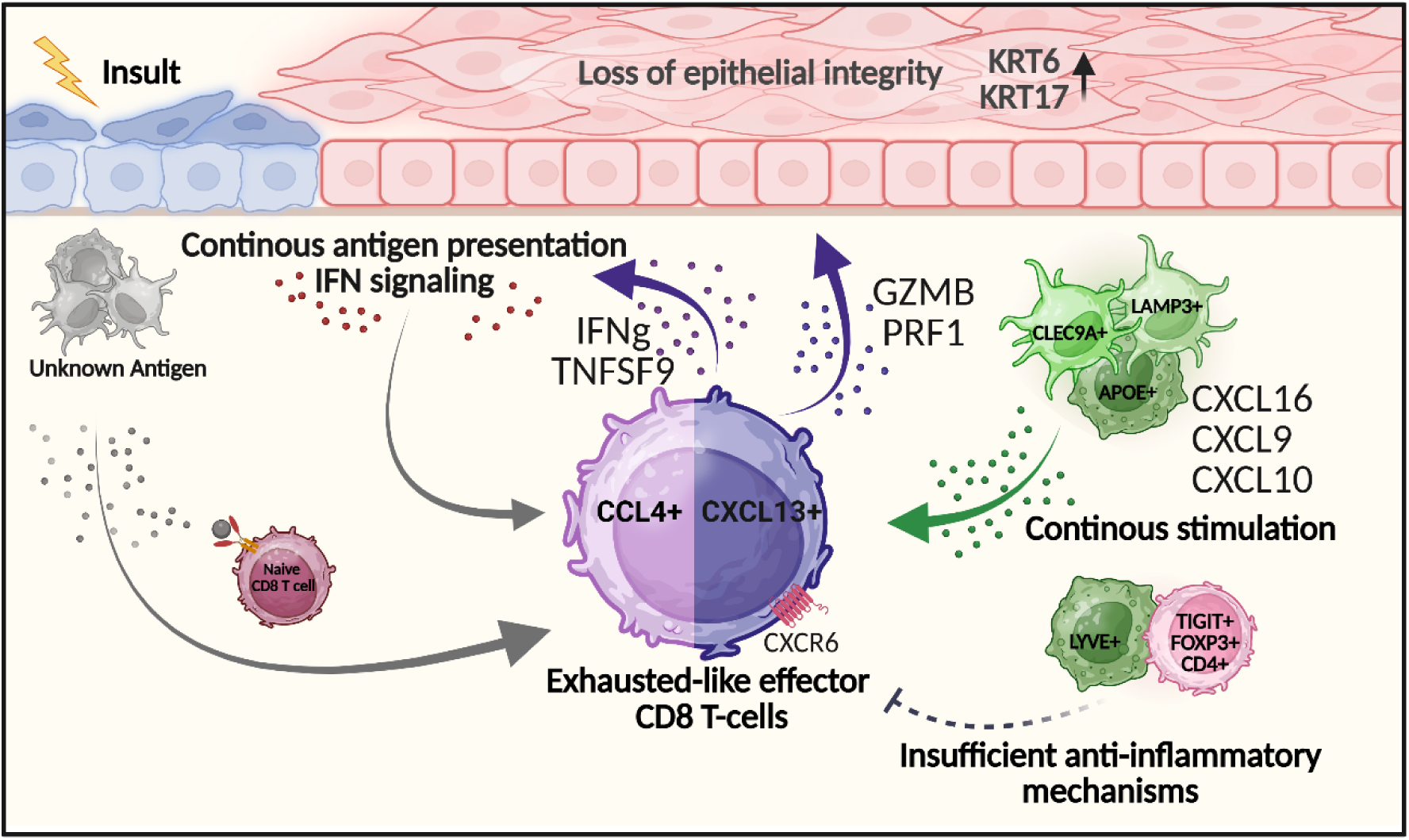
A mechanistic model summarizes the progression from initial epithelial barrier disruption and persistent antigen exposure to the generation of exhausted-like CD8⁺ T cells. These cells, marked by CCL4 and CXCL13 expression, retain cytotoxic potential and contribute to chronic tissue injury indicated by the expression of damage-associated keratins (KRT6 and KRT17). Failed anti-inflammatory regulation and sustained recruitment via CXCL16–CXCR6 / CXCL9/CXCL10-CXCR3 interactions support the maintenance of pathogenic CD8⁺T cell populations. Antigen-presenting cells (e.g., CLEC9A⁺ and AIRE⁺ cells) may perpetuate activation, creating a feedback loop of immune-driven tissue damage.

## DISCUSSION

Despite ongoing advances in our understanding of cGVHD pathophysiology, it remains a significant clinical challenge following alloHCT. Studies focusing on peripheral blood samples and murine models have provided insight into cGVHD central pathways. However, direct characterization of tissue-infiltrating cells in patients remains limited, and the underlying causes of disease onset in specific patient populations and target organs remain a subject of ongoing debate. In this study, we employed scRNA-seq and multiplex IHC to dissect the cellular and molecular landscape of OM in patients with cGVHD. Our findings reveal marked inter-patient heterogeneity and complexity of AG status and uncover common dysregulated non-immune cell populations that may contribute to sustained antigen presentation and chronic inflammation in GV patient OM. Notably, we provide comprehensive characterization of the immune cell infiltrate in affected OM and closely identify T_EXEF_ cells as a key population implicated in localized tissue damage and clinical cGVHD severity.

Several studies have highlighted the involvement of non-immune cells and APCs in GVHD pathogenesis in other organs (18, 28–30) and oral mucosa (10). Preclinical models have shown that epithelial and endothelial cells upregulate pattern recognition receptors (PRRs) and MHC class II molecules in response to inflammatory cues, enabling them to act as non-traditional APCs. Notably, selective blockade of co-stimulatory molecules on these cell types has been shown to attenuate GVHD initiation (19, 20), reinforcing their role in sustaining immune activation. In the present study, oral mucosal epithelia, endothelia, and fibroblasts exhibited high expression of genes associated with antigen presentation and immune cell recruitment. Epithelial cells, in particular, had increased expression of stress- and damage-associated keratins—KRT6A, KRT16, and KRT17—that were previously implicated in tissue inflammation and remodeling (25, 26). Among these, KRT17 is particularly relevant, as its dysregulated expression has been shown to promote the secretion of pro-inflammatory cytokines such as IFNG and IL17A (25) which are both central to cGVHD pathogenesis. Notably, cell–cell interaction analysis using LIANA further revealed a marked reduction in ECM interactions, including FN1-SDC1, in cGVHD patients, suggesting impaired tissue integrity and increased damage. This disruption likely reflects enhanced matrix degradation or impaired ECM synthesis, contributing to increased release of damage-associated molecular patterns (DAMPs) and sustained inflammatory signaling. Together, these data suggest that non-immune cells contribute not only as targets of inflammation but as active amplifiers of immune dysregulation in the oral mucosa.

In addition to changes in epithelial and stromal compartments, we observed significant alterations in myeloid cell polarization and function in cGVHD patients. Macrophages and DCs are highly responsive to tissue-derived signals, particularly those released by stressed or damaged non-immune cells (31). In this context, DAMPs act as potent activators of myeloid cells, amplifying local inflammation through engagement of pattern recognition receptors (32). Consistent with this, macrophages in the oral mucosa of cGVHD patients showed a pronounced shift toward an M1-like pro-inflammatory phenotype, alongside a relative reduction in M2-like macrophages that are typically associated with tissue repair and immune resolution (29, 30). This polarization imbalance was accompanied by increased scores for antigen presentation and phagocytic activity in both APCs and non-immune cells, reflecting a mucosal environment primed for sustained immune activation. Furthermore, the observed disruption in ECM interactions likely exacerbates this inflammatory state by increasing the availability of DAMPs, thereby reinforcing APC activation and perpetuating the inflammatory cascade.

Building on this pro-inflammatory milieu, our findings point to a critical role for dysregulated CD8⁺ T cells, particularly exhausted-like subsets, in perpetuating tissue damage in cGVHD. While donor naïve CD4+ T-cells are an important subset involved in cGVHD development (13–15), recent preclinical studies have identified functionally exhausted, yet still cytotoxic, CD8⁺ T cells as contributors to disease pathology through cytokine secretion and tissue cytotoxicity (33–35). These exhausted-like cells are unlikely to respond to regulatory signals to cease cytotoxic activity. The contribution of T-cell exhaustion to the development and persistence of alloimmunity remains underexplored in cGVHD. In our dataset, we observed an expansion of two distinct T_EXEF_ populations, CD8T_4 and CD8T_5, characterized by features of both effector function and exhaustion. More notably, these populations appear to follow a differentiation trajectory marked by a shift in effector profile. CD8T_4 cells express high levels of CCL4 and TNFSF9, indicative of a transitional or pre-exhausted state capable of cytokine production and immune activation. In contrast, CD8T_5 cells exhibit increased expression of CXCL13 and granzyme B, suggestive of terminal exhaustion with retained cytotoxic potential. Although CXCL13 is classically associated with B-cell recruitment (36, 37), we did not detect B cells in our scRNA-seq cohort, consistent with prior studies showing that tertiary lymphoid structures are rare in oral cGVHD (7, 9, 38). However, emerging evidence suggests that CXCL13 can also recruit CXCR3⁺ effector T cells (39), potentially reinforcing a self-sustaining cytotoxic loop within the inflamed mucosa. This T_EXEF_ cell axis may be further amplified by DAMP signaling from damaged stromal cells, linking epithelial stress to chronic immune activation. Cell–cell interaction analysis supported these findings, revealing strong communication between myeloid cells, non-immune stromal cells, and CD8⁺ T cells via integrin and chemokine signaling pathways. Notably, the CXCL16–CXCR6 axis was uniquely enriched between macrophages and the terminally exhausted CD8T_5 cluster. CXCL16–CXCR6 signaling has been implicated in the retention of CD8⁺ T cells at sites of chronic antigen exposure and contributes to persistent inflammation in autoimmune diseases (40). Critically, expression of CCL4 or expression of CXCL13 within the CD8^+^T cell subset were each positively correlated with clinical severity of oral cGVHD in these patients which suggests a clinically relevant role for these T_EXEF_ cells.

Despite the close proximity of FOXP3⁺ regulatory T cells to cytotoxic T cells, their presence may be insufficient to restore immune balance in this inflammatory niche, highlighting a potential breakdown of local immune regulation in oral cGVHD. These findings suggest that anti-inflammatory mechanisms remain insufficient to counterbalance the dominant pro-inflammatory signals in cGVHD. This may reflect a functional impairment of regulatory T cells or an inability to suppress highly cytotoxic, T_EXEF_ cells that persist in the tissue. Once CD8⁺ T cells have transitioned into a terminally exhausted or cytolytic state, their regulatory reprogramming possibly becomes limited—thereby locking the tissue into a state of sustained damage. Similar shifts in exhaustion phenotype have been described in other chronic inflammatory diseases and cancer (41), suggesting this may represent a broader mechanism of immune escape or persistence that contributes to tissue pathology in cGVHD. These findings underscore the need to better understand the plasticity, function, and regulation of exhausted CD8⁺ T cells in this context, especially as potential therapeutic targets

A pivotal yet underexplored concept in this context is the threshold model of immune activation, which hypothesizes that regulatory circuits enforce a signaling threshold that must be surpassed to trigger inflammation—an idea well established in cancer and autoimmune biology (42). In cGVHD, this model suggests that initial tissue injury may tip the balance toward chronic inflammation if compensatory regulatory mechanisms fail. This is especially relevant in transplanted patients, where ongoing low-level immune activation may push tissues into a para-inflammatory state, an intermediate between homeostasis and full-blown inflammation (43). Importantly, this threshold is not fixed, and it varies by tissue, individual genetics, and local cellular context (44–46). In OM, our single-cell analysis highlights a unique tissue-specific immune landscape shaped by stromal-immune crosstalk and CD8⁺ T cell dysfunction. Whether similar T_EXEF_ cell states exist in other oral tissues affected by cGVHD, such as the salivary glands, remains an open question, but one with significant implications for understanding disease heterogeneity and therapeutic targeting.

This single site study has inherent limitations, including the small numbers of patient samples included in each group from which high-quality data suitable for deep analysis could be derived. Additional work is needed to ensure that these findings are generalizable to new-onset cGVHD patients at other transplant centers and to determine if these findings can be extended to mucosal sites beyond the oral cavity.

In this study, we have identified overall changes in the cellular landscape and key dysregulated cell populations in OM GV, laying the groundwork for further work in several key areas. Functional validation of the T_EXEF_ cell subsets we identified is crucial to determine their true effector potential, plasticity, and response to checkpoint or cytokine modulation. Spatial and temporal profiling, especially with technologies such as spatial transcriptomics or multiplexed imaging, will clarify whether these cells occupy distinct niches or interact dynamically with myeloid or stromal populations. Also, studying the molecular triggers that shift T cells from an early activation state to exhaustion in tissue may uncover actionable checkpoints to prevent irreversible damage. This investigation implicates T_EXEF_ cells in OM GV as potential therapeutic targets to improve treatment for cGVHD and restore immune homeostasis. Future work to define how the immune threshold varies across patients and tissues could improve risk stratification and inform tissue-specific therapies for oral cGVHD.

## MATERIALS AND METHODS

### Study design

This study was designed to prospectively characterize the cellular profile of oral buccal mucosa (OM) in patients and identify cell types and pathways associated with chronic graft-versus-host disease (cGVHD). To achieve this, we conducted single-cell RNA sequencing (scRNA-seq) and flow cytometric analysis on OM biopsies obtained from patients at time- or event-driven timepoints. Multiplex immunostaining was further performed to confirm the presence of specific cell types. The number of patients used per methodology and the statistical tests performed are listed in the respective figure legends.

### Patient enrollment and tissue collection

Patients were sampled, including OM biopsy, at the NIH Clinical Center following informed consent and enrollment on IRB-approved protocols registered at clinicaltrials.gov (NCT00092235 and NCT03602599) conducted in accordance with the Declaration of Helsinki. Clinical assessment including Oral Mucositis Rating Scale (OMRS) scoring was completed by a calibrated dentist in the NIH Dental Clinic. Clinical pathology was scored by NIH Clinical Center Pathology based on the NIH Consensus Criteria for histopathologic diagnosis of oral cGVHD (17). Both sexes were included in the cohort; however, due to sequential patient enrollment, the sex distribution is not balanced across groups, and no sex-specific sub analyses were conducted. Full demographic details, including age and sex and sample size, for all patient cohorts, are detailed in Table S1.

A 6mm punch biopsy was taken from the oral mucosa from affected or clinical normal tissue from NG (*n=3*), AG (*n=4*) and GV (*n=9*) patients at predetermined time or event-driven points after alloHCT. The tissue was sectioned consistently such that part was used for scRNA-seq or FACS, one part was formalin-fixed and sent to Pathology, and one part was further sectioned then embedded in OCT or RNAlater.

### Histology and Hematoxylin and Eosin Staining

Formalin fixed, paraffin embedded (FFPE) OM sections were processed and stained with Hematoxylin and Eosin (H&E) by NIH Clinical Pathology and then read by a pathologist. H&E sections were imaged at brightfield using a NanoZoomer S60 (Hamamatsu Photonics) on 40X objective lens and images were exported using NDP.view2.

### Tissue dissociation and processing

Fresh OM tissues were dissociated mechanically and then enzymatically with Collagenase P (Worthington Biochemical Corporation) and DNase (Sigma). They were further processed through the gentleMACS Dissociator (Miltenyi) and incubated at 37°C with shaker for 1 hr. Tissue lysate was again dissociated via gentleMACS Dissociator and was filtered through a 70 μm strainer (Miltenyi), washed in culture medium [RPMI-1640 (Gibco) with 2% FBS], and resuspended in RBC lysis buffer for 1 min. Cells were centrifuged (10 mins at 300*g*, 4°C), washed, and resuspended in culture medium. Cell counts and viability were confirmed using Countess II automated cell counter (Thermo Fisher) and were used for downstream scRNA-seq and flow cytometry analysis.

### Single-cell mRNA sequencing

#### Library construction and sequencing

Single-cell suspensions were loaded onto a 10X Chromium Controller (10X Genomics), and library preparation was performed according to the manufacturer’s instructions for the 10X Chromium Next GEM Single Cell Library kit v3 (10X Genomics). Quantification of cDNA was performed using an Agilent Bioanalyzer 2100 (Agilent technologies, Santa Clara, CA). Libraries were then sequenced on a NextSeq 2000 sequencer (Illumina) using 10X Genomics recommended reads configuration.

Details regarding the initial number of captured cells per biopsy, average reads per cell, and other sequencing metrics are provided in Table S2. Doublets, unclassified cells, or clusters with high mitochondrial content were excluded from further analysis.

#### scRNA-seq data processing

Cell Ranger files from 10X were imported to SEURAT v4 using R v.4.4.0 and processed for clustering using default pipeline (47)(SEURAT v4). For quality control, cells with less than 200 genes and with >10-25 % of UMIs mapping to mitochondrial genes were excluded from the subsequent analyses. Data was normalized using SCT transformation before harmony integration (48) all 15 patients’ datasets. Principal component analysis (PCA) was performed and the first 30 PCs were retained. ‘Clustree’ package (49) was used to determine an optimal resolution for clustering. ‘FindAllMarkers’ function in Seurat was used to identify specific genes for each cell cluster. (Data file S1). Differentially expressed genes were performed using MAST (50) and were defined by thresholds of log2FoldChange > 0 and pAdjustMethod (p. adj) < 0.05. We applied clusterProfiler R package (51) (version 4.0) for functional enrichment analysis using the terms from Gene Ontology Biological Process (GO BP).

#### Gene signature score

Functional states of cell populations were evaluated by calculating gene signature scores using the UCell R package (52) (v2.8). For non-immune cells and macrophages, the antigen presentation score was derived from the GOBP_ANTIGEN_PROCESSING_AND_PRESENTATION gene set (GO:0019882). Chemotaxis and inflammatory response regulation scores were calculated using GOBP_LEUKOCYTE_CHEMOTAXIS (GO:0030595) and GOBP_REGULATION_OF_INFLAMMATORY_RESPONSE (GO:0050727), respectively (Table S3). M1 and M2 polarization scores were determined based on gene sets from published studies (27), while phagocytosis scores were defined using GOBP_POSITIVE_REGULATION_OF_PHAGOCYTOSIS (GO:0050766) (Table S4). For T cells, residency, cytotoxic, and exhaustion scores were computed using established markers and gene sets from prior studies (53–55) (Table S5). Scores were calculated with the “area under the curve” (AUC) method, using AUCell R package. Scores were compared across clusters and visualized using violin plots and UMAP projections.

#### Trajectory and pseudotime analysis

Trajectory analysis of CD8 T cells was performed using Slingshot (v2.2.0) (56) via the SeuratExtend R package. The post-processed Seurat object was converted to a single cell experiment object. Slingshot was applied with sample names as cluster identifiers, designating CD8T_3 as the starting cluster for pseudotime ordering. Trajectories were visualized using plotGenePseudotime.

### Cell-cell communication analysis

Cell-cell communication was analyzed using CellChat (v1.6.1) (http://www.cellchat.org/). The normalized gene expression matrix and metadata (with sample names as cluster identifiers) were extracted from the Seurat object and converted into a CellChat object. The human CellChatDB was used to identify ligand-receptor interactions. Probabilities of communication were computed using computeCommunProb with default parameters, and significant interactions were filtered at a p-value threshold of 0.05. Communication networks were visualized using netVisual_circle and netVisual_heatmap. Ligand–receptor interaction analysis was also conducted using LIANA via *consensus* pipeline (57) (LIANA-py; version 1.5.1) and tensor-cell2cell approach (58) was used to assess patterns of communications across samples.

### Immunohistochemistry (IHC)

FFPE sections were deparaffinized, dewaxed in xylene substitute (Sigma) and rehydrated using a decreasing sequence of graded ethanols. Antigen retrieval with Tris-EDTA buffer (pH 9) was performed using a pressure cooker for 10 min and blocked in 5% donkey serum (Jackson Immunoresearch) for 30 min at RT. Slides were incubated with primary antibodies in PBS_0.1%_Bovine Serum Albumin (BSA) overnight at 4°C. The next day, unconjugated antibodies were incubated with secondary antibodies for 30 minutes at RT, followed by incubation with conjugated primary antibody for 2 hours at RT in dark conditions prior to 4′,6-diamidino-2-phenylindole (DAPI; Thermo Scientific; 1:2000) incubation for 5 minutes. To mount, slides were dried and then mounted using Fluoro-gel (EMS). Tissues were imaged using a Leica SP8 WLL inverted confocal microscope at 40× magnification. A list of antibodies used for IHC is available on Table S6.

### IHC quantification analysis

For CD45 staining, whole OM sections were imaged on the Nikon TiE#2 Widefield microscope at 20x magnification and was quantified using a machine learning pixel classification tool in Imaris 9.9.0, in conjunction with Labkit pixel training and segmentation in FIJI software. Cell counts were normalized by the area of the tissue (µm^2^).

For CD3, CD8, CCL4 and CXCL13 panel, tile scan images were captured at a magnification of 40x using LEICA SP8 (with STED and DLS) microscope. Images were processed in Fiji and were quantified using QuPath (v.0.5.1) and normalized to area.

### Flow Cytometry

Single-cell suspensions from OM tissues were incubated with Live/Dead fixable dye (Zombie UV, Invitrogen) to exclude dead cells and blocked with TruStain FcX (Biolegend, San Diego, CA). Cells were then incubated with conjugated antibodies anti CD45, CD3, CD8 and CD4 for 30 min at 4°C in dark, followed by fixation with paraformaldehyde. Cells were acquired on BD LSRFortessa using FACSDiVa software (BD Biosciences, San Jose, CA). Antibodies and reagents used are listed in Table S6. Data was analyzed with FlowJo software (Version 10.9, Ashland, OR: Becton, Dickinson and Co.).

### CODEX staining and Analysis

The CODEX staining was performed on square (22 × 22 mm) glass coverslips (72204-10, Electron Microscopy Sciences, Hatfield, PA) pre-treated with Poly L-Lysine (Sigma, St. Louis, MO) according to Akoya Biosciences. We stained for CD8, TOX, LAG3, TIGIT, IFNG, FoxP3 and TIM3. Antibodies and reagents used are listed in Table S6. Tissue sections were stained with an antibody cocktail consisting of 2-5 μl of each antibody per tissue. Raw TIFF images produced during image acquisition were processed using the CODEX image processer (version 1.8.3.14). The output of this image processing was tiled images corresponding to all fluorescence channels and imaging cycles that were then visualized and analyzed using HALO software (Version 3.3.2541.383, Indica Labs Inc., Albuquerque, NM). Segmentation of cells was performed using the nuclear channel and the cell cytoplasm was defined as a fixed width ring around each nucleus. Nuclear segmentation settings were optimized by visual verification of segmentation performance on random subsets of cells aiming to minimize the number of over segmentations, under segmentation, detected artefacts and missed cells. All nucleated cells were first identified by positive nuclear signals and the cell phenotypes were defined based on the combination if signals described in using HALO and/or positive and negative criteria.

### Statistical analysis

Survival analysis, one-way ANOVA with Tukey’s multiple comparisons test or Mann-Whitney post-hoc testing was used to assess statistical significance using GraphPad Prism software (Version 10.0.2, La Jolla, CA). Significance is reported as *p ≤ 0.05, **p ≤ 0.01, ***p ≤ 0.001, ****p ≤ 0.0001.

### Software

R version 4.3.2 was used for the analyses and figure creation. Python 3 was used for cell-cell interaction. Illustrations were created with BioRender.com and Adobe Photoshop.

## Data Availability

Single-cell RNA-seq data (will be) deposited at GEO and (will be) publicly available as of the date of publication (Accession #). Microscopy data reported in this paper will be shared by the lead contact upon request. This manuscript does not report original code. All codes were used in this study in alignment with recommendations made by authors of R packages in their respective user guide. Any additional information required to reanalyze the data reported in this paper is available from the lead contact upon request.

## ACKNOWLEDGEMENTS

This study was funded by the Intramural Program of the National Institute of Dental and Craniofacial Research, National Institutes of Health, Bethesda, MD, USA (ZIA DE000747 to JWM). Thank you to the patients of the NIH Clinical Center and their families, without whom this work would not be possible. The authors thank the NIDCR Combined Technical Core (ZIC DE000729), the NIDCR Imaging Core (ZIC DE000750-01), and the NIDCD/NIDCR Genomics and Computational Biology Core (ZIC DC000086) for excellent technical assistance and guidance. Thank you to Licia Masuch and Pam Orzechowski for their clinical trial data and regulatory management.

## AUTHOR CONTRIBUTIONS

ACC, RS, MHA, and JWM wrote the manuscript. ACC, RS, MHA, DM, NK, DFK, AJ, FAB, DPM, JTD, CHK, DEK, and JWM analyzed specimens and data. JWM, JTN, SMG, MKH, DEK, CGK, SZP provided patient care and acquisition of clinical specimens. All authors contributed to the critical revision of the manuscript and approved the final submitted version.

## SUPPLEMENTARY MATERIALS

Figs. S1 to S6

Tables S1 to S6

## REFERENCES

1. Flowers, M. E., and Martin, P. J. (2015) How we treat chronic graft-versus-host disease Blood 125, 606–615

2. Carpenter, P. A., Gooley, T. A., Boiko, J., Lee, C. J., Burroughs, L. M., Mehta, R. et al. (2024) Decreasing chronic graft-versus-host disease rates in all populations Blood Adv 8, 5829–5837

3. Bachier, C. R., Aggarwal, S. K., Hennegan, K., Milgroom, A., Francis, K., Dehipawala, S. et al. (2021) Epidemiology and Treatment of Chronic Graft-versus-Host Disease Post-Allogeneic Hematopoietic Cell Transplantation: A US Claims Analysis Transplant Cell Ther 27, 504 e501–504 e506

4. Curtis, D. J., Patil, S. S., Reynolds, J., Purtill, D., Lewis, C., Ritchie, D. S. et al. (2025) Graft-versus-Host Disease Prophylaxis with Cyclophosphamide and Cyclosporin N Engl J Med 393, 243–254

5. Dean, D., and Sroussi, H. (2022) Oral Chronic Graft-Versus-Host Disease Front Oral Health 3, 903154

6. Johnson, L. B., Oh, U., Rothen, M., Sroussi, H. Y., Dean, D. R., Lloid, C. M. et al. (2022) A Review of Oral Chronic Graft-Versus-Host Disease: Considerations for dental hygiene practice J Dent Hyg 96, 6–17

7. Fall-Dickson, J. M., Pavletic, S. Z., Mays, J. W., and Schubert, M. M. (2019) Oral Complications of Chronic Graft-Versus-Host Disease J Natl Cancer Inst Monogr 2019,

8. Cooke, K. R., Luznik, L., Sarantopoulos, S., Hakim, F. T., Jagasia, M., Fowler, D. H. et al. (2017) The Biology of Chronic Graft-versus-Host Disease: A Task Force Report from the National Institutes of Health Consensus Development Project on Criteria for Clinical Trials in Chronic Graft-versus-Host Disease Biol Blood Marrow Transplant 23, 211–234

9. Nguyen, J. T., Jessri, M., Costa-da-Silva, A. C., Sharma, R., Mays, J. W., and Treister, N. S. (2024) Oral Chronic Graft-Versus-Host Disease: Pathogenesis, Diagnosis, Current Treatment, and Emerging Therapies Int J Mol Sci 25,

10. Sharma, R., Holtzman, N. G., Pusic, I., Cutler, C., Treister, N., Mehta, R. S. et al. (2025) Belumosudil reduces oral chronic graft-versus-host disease tissue inflammation and fibrosis: a ROCKstar companion study Blood Adv 9, 3479–3494

11. Wu, T., Young, J. S., Johnston, H., Ni, X., Deng, R., Racine, J. et al. (2013) Thymic damage, impaired negative selection, and development of chronic graft-versus-host disease caused by donor CD4+ and CD8+ T cells J Immunol 191, 488–499

12. Zhang, X., He, J., Zhao, K., Liu, S., Xuan, L., Chen, S. et al. (2023) Mesenchymal stromal cells ameliorate chronic GVHD by boosting thymic regeneration in a CCR9-dependent manner in mice Blood Adv 7, 5359–5373

13. Chen, B. J., Deoliveira, D., Cui, X., Le, N. T., Son, J., Whitesides, J. F. et al. (2007) Inability of memory T cells to induce graft-versus-host disease is a result of an abortive alloresponse Blood 109, 3115–3123

14. Dutt, S., Tseng, D., Ermann, J., George, T. I., Liu, Y. P., Davis, C. R. et al. (2007) Naive and memory T cells induce different types of graft-versus-host disease J Immunol 179, 6547–6554

15. Zheng, H., Matte-Martone, C., Jain, D., McNiff, J., and Shlomchik, W. D. (2009) Central memory CD8+ T cells induce graft-versus-host disease and mediate graft-versus-leukemia J Immunol 182, 5938–5948

16. Bleakley, M., Sehgal, A., Seropian, S., Biernacki, M. A., Krakow, E. F., Dahlberg, A. et al. (2022) Naive T-Cell Depletion to Prevent Chronic Graft-Versus-Host Disease J Clin Oncol 40, 1174–1185

17. Jagasia, M. H., Greinix, H. T., Arora, M., Williams, K. M., Wolff, D., Cowen, E. W. et al. (2015) National Institutes of Health Consensus Development Project on Criteria for Clinical Trials in Chronic Graft-versus-Host Disease: I. The 2014 Diagnosis and Staging Working Group report Biol Blood Marrow Transplant 21, 389–401 e381

18. Dainichi, T., Kabashima, K., Ivanov, II, and Goto, Y. (2021) Editorial: Regulation of Immunity by Non-Immune Cells Front Immunol 12, 770847

19. Neidemire-Colley, L., Robert, J., Ackaoui, A., Dorrance, A. M., Guimond, M., and Ranganathan, P. (2022) Role of endothelial cells in graft-versus-host disease Front Immunol 13, 1033490

20. Koyama, M., Mukhopadhyay, P., Schuster, I. S., Henden, A. S., Hulsdunker, J., Varelias, A. et al. (2019) MHC Class II Antigen Presentation by the Intestinal Epithelium Initiates Graft-versus-Host Disease and Is Influenced by the Microbiota Immunity 51, 885–898 e887

21. Zima, K., Purzycka-Bohdan, D., Szczerkowska-Dobosz, A., and Gabig-Ciminska, M. (2024) Keratinocyte-Mediated Antigen Presentation in Psoriasis: Preliminary Insights from In Vitro Studies Int J Mol Sci 25,

22. Imanguli, M. M., Swaim, W. D., League, S. C., Gress, R. E., Pavletic, S. Z., and Hakim, F. T. (2009) Increased T-bet+ cytotoxic effectors and type I interferon-mediated processes in chronic graft-versus-host disease of the oral mucosa Blood 113, 3620–3630

23. Cohen, E., Johnson, C. N., Wasikowski, R., Billi, A. C., Tsoi, L. C., Kahlenberg, J. M. et al. (2024) Significance of stress keratin expression in normal and diseased epithelia iScience 27, 108805

24. Wang, T. H., Shen, Y. W., Chen, H. Y., Chen, C. C., Lin, N. C., Shih, Y. H. et al. (2024) Arecoline Induces ROS Accumulation, Transcription of Proinflammatory Factors, and Expression of KRT6 in Oral Epithelial Cells Biomedicines 12,

25. Romashin, D. D., Tolstova, T. V., Varshaver, A. M., Kozhin, P. M., Rusanov, A. L., and Luzgina, N. G. (2024) Keratins 6, 16, and 17 in Health and Disease: A Summary of Recent Findings Curr Issues Mol Biol 46, 8627–8641

26. Jansen, S. A., Nieuwenhuis, E. E. S., Hanash, A. M., and Lindemans, C. A. (2022) Challenges and opportunities targeting mechanisms of epithelial injury and recovery in acute intestinal graft-versus-host disease Mucosal Immunol 15, 605–619

27. Guimaraes, G. R., Maklouf, G. R., Teixeira, C. E., de Oliveira Santos, L., Tessarollo, N. G., de Toledo, N. E. et al. (2024) Single-cell resolution characterization of myeloid-derived cell states with implication in cancer outcome Nat Commun 15, 5694

28. Du, J., Paz, K., Flynn, R., Vulic, A., Robinson, T. M., Lineburg, K. E. et al. (2017) Pirfenidone ameliorates murine chronic GVHD through inhibition of macrophage infiltration and TGF-beta production Blood 129, 2570–2580

29. Alexander, K. A., Flynn, R., Lineburg, K. E., Kuns, R. D., Teal, B. E., Olver, S. D. et al. (2014) CSF-1-dependant donor-derived macrophages mediate chronic graft-versus-host disease J Clin Invest 124, 4266–4280

30. Hong, Y. Q., Wan, B., and Li, X. F. (2020) Macrophage regulation of graft-vs-host disease World J Clin Cases 8, 1793–1805

31. Del Fresno, C., and Sancho, D. (2021) Myeloid cells in sensing of tissue damage Curr Opin Immunol 68, 34–40

32. Dwyer, G. K., and Turnquist, H. R. (2021) Untangling Local Pro-Inflammatory, Reparative, and Regulatory Damage-Associated Molecular-Patterns (DAMPs) Pathways to Improve Transplant Outcomes Front Immunol 12, 611910

33. Kosugi-Kanaya, M., Ueha, S., Abe, J., Shichino, S., Shand, F. H. W., Morikawa, T. et al. (2017) Long-Lasting Graft-Derived Donor T Cells Contribute to the Pathogenesis of Chronic Graft-versus-Host Disease in Mice Front Immunol 8, 1842

34. Gao, Y., Liu, R., Shi, J., Shan, W., Zhou, H., Chen, Z. et al. (2025) Clonal GZMK(+)CD8(+) T cells are identified as a hallmark of the pathogenesis of cGVHD-induced bronchiolitis obliterans syndrome after allogeneic hematopoietic stem cell transplantation EBioMedicine 112, 105535

35. Mellors, P. W., Lange, A. N., Casino Remondo, B., Shestov, M., Planer, J. D., Peterson, A. R. et al. (2025) Shared roles of immune and stromal cells in the pathogenesis of human bronchiolitis obliterans syndrome JCI Insight 10,

36. Gunn, M. D., Ngo, V. N., Ansel, K. M., Ekland, E. H., Cyster, J. G., and Williams, L. T. (1998) A B-cell-homing chemokine made in lymphoid follicles activates Burkitt’s lymphoma receptor-1 Nature 391, 799–803

37. Legler, D. F., Loetscher, M., Roos, R. S., Clark-Lewis, I., Baggiolini, M., and Moser, B. (1998) B cell-attracting chemokine 1, a human CXC chemokine expressed in lymphoid tissues, selectively attracts B lymphocytes via BLR1/CXCR5 J Exp Med 187, 655–660

38. Tollemar, V., Garming Legert, K., and Sugars, R. V. (2023) Perspectives on oral chronic graft-versus-host disease from immunobiology to morbid diagnoses Front Immunol 14, 1151493

39. Jenh, C. H., Cox, M. A., Hipkin, W., Lu, T., Pugliese-Sivo, C., Gonsiorek, W. et al. (2001) Human B cell-attracting chemokine 1 (BCA-1; CXCL13) is an agonist for the human CXCR3 receptor Cytokine 15, 113–121

40. Bao, N., Fu, B., Zhong, X., Jia, S., Ren, Z., Wang, H. et al. (2023) Role of the CXCR6/CXCL16 axis in autoimmune diseases Int Immunopharmacol 121, 110530

41. Tietscher, S., Wagner, J., Anzeneder, T., Langwieder, C., Rees, M., Sobottka, B. et al. (2023) A comprehensive single-cell map of T cell exhaustion-associated immune environments in human breast cancer Nat Commun 14, 98

42. Guram, K., Kim, S. S., Wu, V., Sanders, P. D., Patel, S., Schoenberger, S. P. et al. (2019) A Threshold Model for T-Cell Activation in the Era of Checkpoint Blockade Immunotherapy Front Immunol 10, 491

43. Medzhitov, R. (2008) Origin and physiological roles of inflammation Nature 454, 428–435

44. Oyesola, O., Downie, A. E., Howard, N., Barre, R. S., Kiwanuka, K., Zaldana, K. et al. (2023) Genetic and Environmental interactions contribute to immune variation in rewilded mice bioRxiv 10.1101/2023.03.17.533121

45. Mangino, M., Roederer, M., Beddall, M. H., Nestle, F. O., and Spector, T. D. (2017) Innate and adaptive immune traits are differentially affected by genetic and environmental factors Nat Commun 8, 13850

46. Liston, A., Humblet-Baron, S., Duffy, D., and Goris, A. (2021) Human immune diversity: from evolution to modernity Nat Immunol 22, 1479–1489

47. Hao, Y., Hao, S., Andersen-Nissen, E., Mauck, W. M., 3rd, Zheng, S., Butler, A. et al. (2021) Integrated analysis of multimodal single-cell data Cell 184, 3573–3587 e3529

48. Korsunsky, I., Millard, N., Fan, J., Slowikowski, K., Zhang, F., Wei, K. et al. (2019) Fast, sensitive and accurate integration of single-cell data with Harmony Nat Methods 16, 1289–1296

49. Zappia, L., and Oshlack, A. (2018) Clustering trees: a visualization for evaluating clusterings at multiple resolutions Gigascience 7,

50. Finak, G., McDavid, A., Yajima, M., Deng, J., Gersuk, V., Shalek, A. K. et al. (2015) MAST: a flexible statistical framework for assessing transcriptional changes and characterizing heterogeneity in single-cell RNA sequencing data Genome Biol 16, 278

51. Yu, G., Wang, L. G., Han, Y., and He, Q. Y. (2012) clusterProfiler: an R package for comparing biological themes among gene clusters OMICS 16, 284–287

52. Andreatta, M., and Carmona, S. J. (2021) UCell: Robust and scalable single-cell gene signature scoring Comput Struct Biotechnol J 19, 3796–3798

53. de Jong, M. J. M., Depuydt, M. A. C., Schaftenaar, F. H., Liu, K., Maters, D., Wezel, A. et al. (2024) Resident Memory T Cells in the Atherosclerotic Lesion Associate With Reduced Macrophage Content and Increased Lesion Stability Arterioscler Thromb Vasc Biol 44, 1318–1329

54. Zhang, J. Y., Wang, X. M., Xing, X., Xu, Z., Zhang, C., Song, J. W. et al. (2020) Single-cell landscape of immunological responses in patients with COVID-19 Nat Immunol 21, 1107–1118

55. Chu, Y., Dai, E., Li, Y., Han, G., Pei, G., Ingram, D. R. et al. (2023) Pan-cancer T cell atlas links a cellular stress response state to immunotherapy resistance Nat Med 29, 1550–1562

56. Street, K., Risso, D., Fletcher, R. B., Das, D., Ngai, J., Yosef, N. et al. (2018) Slingshot: cell lineage and pseudotime inference for single-cell transcriptomics BMC Genomics 19, 477

57. Dimitrov, D., Schafer, P. S. L., Farr, E., Rodriguez-Mier, P., Lobentanzer, S., Badia, I. M. P. et al. (2024) LIANA+ provides an all-in-one framework for cell-cell communication inference Nat Cell Biol 26, 1613–1622

58. Armingol, E., Baghdassarian, H. M., Martino, C., Perez-Lopez, A., Aamodt, C., Knight, R. et al. (2022) Context-aware deconvolution of cell-cell communication with Tensor-cell2cell Nat Commun 13, 3665

