## Supplemental Data for "Exhausted-like effector CD8 T cells mediate immune-stromal interactions at mucosal Chronic Graft-versus-Host Disease onset"

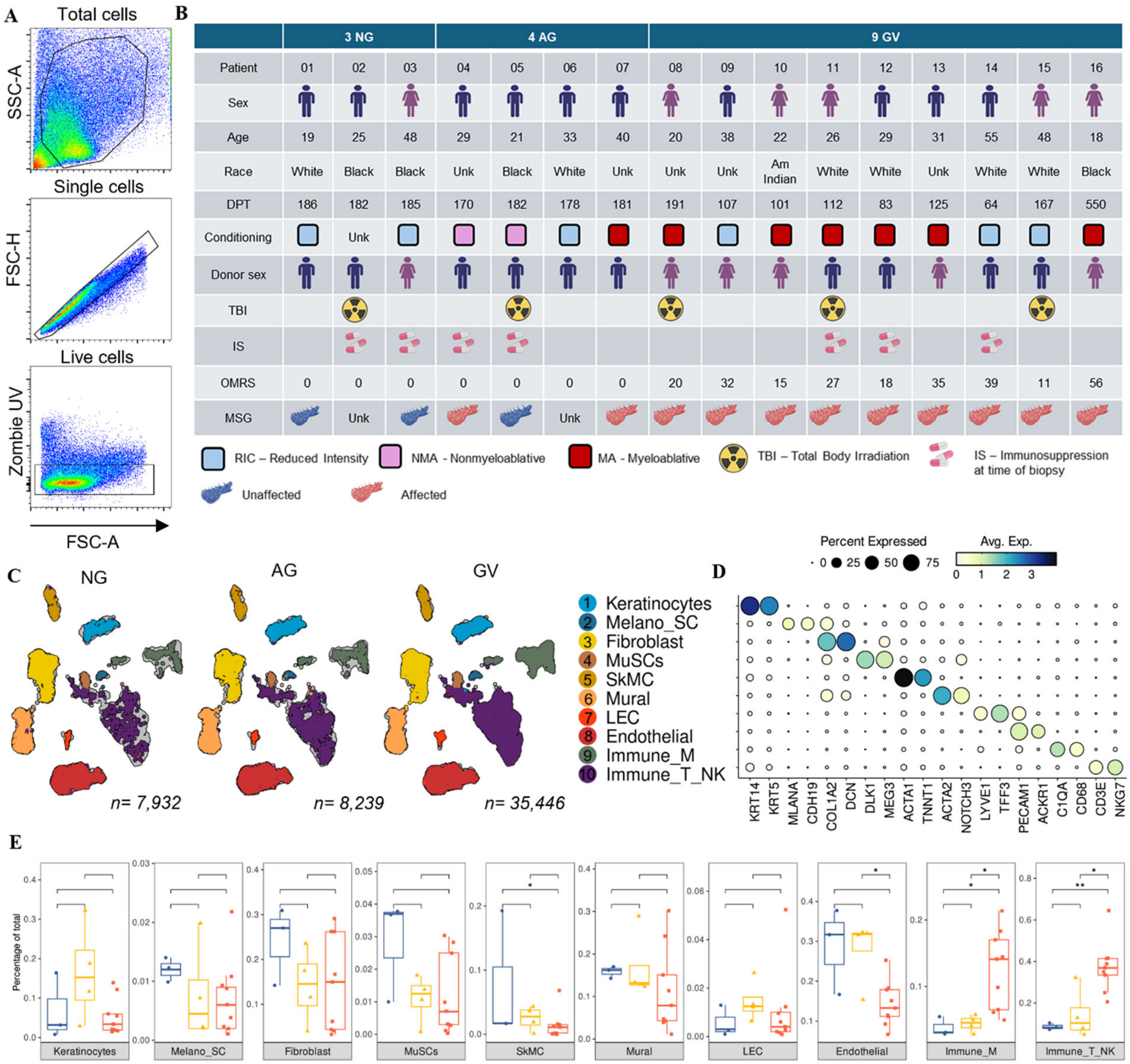

**Supplemental Figure 1: Clinical metadata and cellular composition of oral mucosa in NG, AG, and GV groups. (A)** Flow cytometry gating strategy for isolating viable (Zombie$^{-}$) CD45$^{+}$ immune cells from oral mucosa (OM) biopsies. **(B)** Clinical and demographic characteristics of patients included in the single-cell RNA sequencing analysis. Columns indicate individual patient metadata including gender, age, race, transplant conditioning regimen (RIC: blue, NMA: pink, MA: red), total body irradiation (TBI), donor gender, immunosuppressive treatment (IS), oral cGVHD clinical score (OMRS), and minor salivary gland involvement (MSG). OMRS > 0 indicates clinical oral cGVHD (GV), while AG patients have histological evidence without clinical symptoms. DPT: Days post-transplant. **(C)** UMAP projections of integrated OM single-cell data from NG (n = 7,932 cells), AG (n = 8,239), and GV (n = 35,446) patients, colored by cell type identity (as defined in Fig. 1F). **(D)** Dot plot showing key marker genes used to annotate the 10 identified OM cell types. Dot size indicates the percentage of cells expressing the gene within each cluster; color intensity reflects average expression level. **(E)** Boxplots showing relative frequencies of each cell type across NG (blue), AG (yellow), and GV (red) samples. Significant increases in myeloid and lymphoid populations are observed in GV patients. Each dot represents one patient sample; data shown as mean ± SEM. Statistical comparisons by two-sided Wilcoxon test; $*p \leq 0.05$ considered significant.

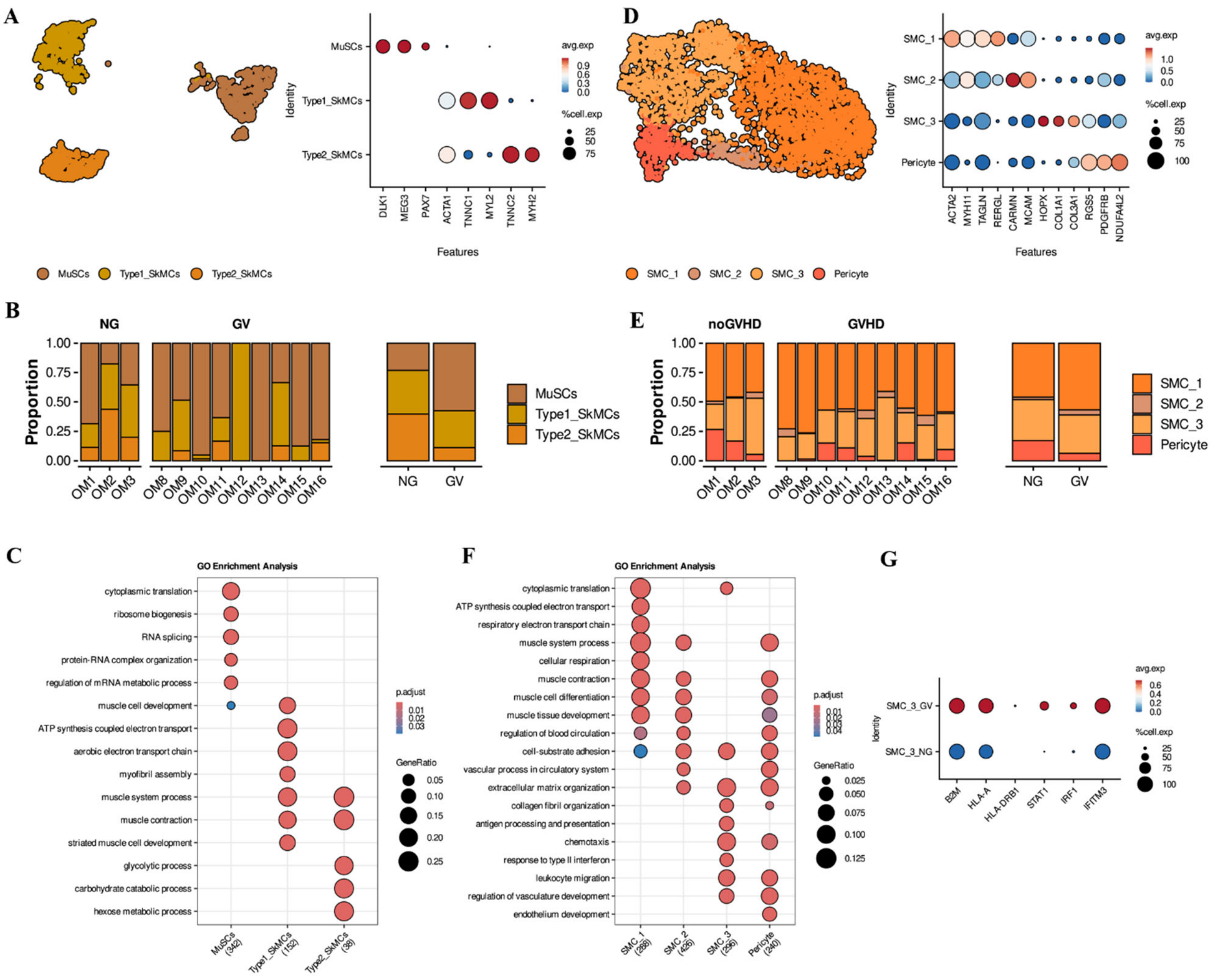

**Supplemental Figure 2. Expanded analysis of non-immune cell subsets: Muscle and Smooth Muscle Cell Alterations in Oral cGVHD. (A)** UMAP (left) of integrated muscle cell populations and dot plot (right) showing expression of representative marker genes for each subcluster **(B)** Bar plots showing the proportion of muscle cell subclusters per individual patient (left) and by group (NG vs. GV, right). Significant differences assessed by two-sided paired Wilcoxon test; $p < 0.05$. Data was assessed by two-sided paired Wilcox test, and p-values < 0.05 was considered statistically significant. **(C)** GO Biological Process (GO BP) pathway enrichment analysis of genes upregulated in each muscle subcluster. Circle size indicates gene ratio; color scale reflects adjusted p-value (Benjamini–Hochberg). **(D)** UMAP (left) and corresponding dot plot (right) of smooth muscle cell (SMC) clusters showing signature gene expression across subpopulations. **(E)** Cell proportion bar plots of SMC subclusters per patient (left) and grouped by NG vs. GV (right). Differences notated as in panel (B). **(F)** GO BP enrichment for genes upregulated in smooth muscle cell subclusters. Significance assessed using one-sided Fisher's exact test with BH correction. **(G)** Dot plot of differentially expressed genes in SMC_3 between NG and GV patients. Dot size reflects percentage of expressing cells; color denotes scaled average expression.

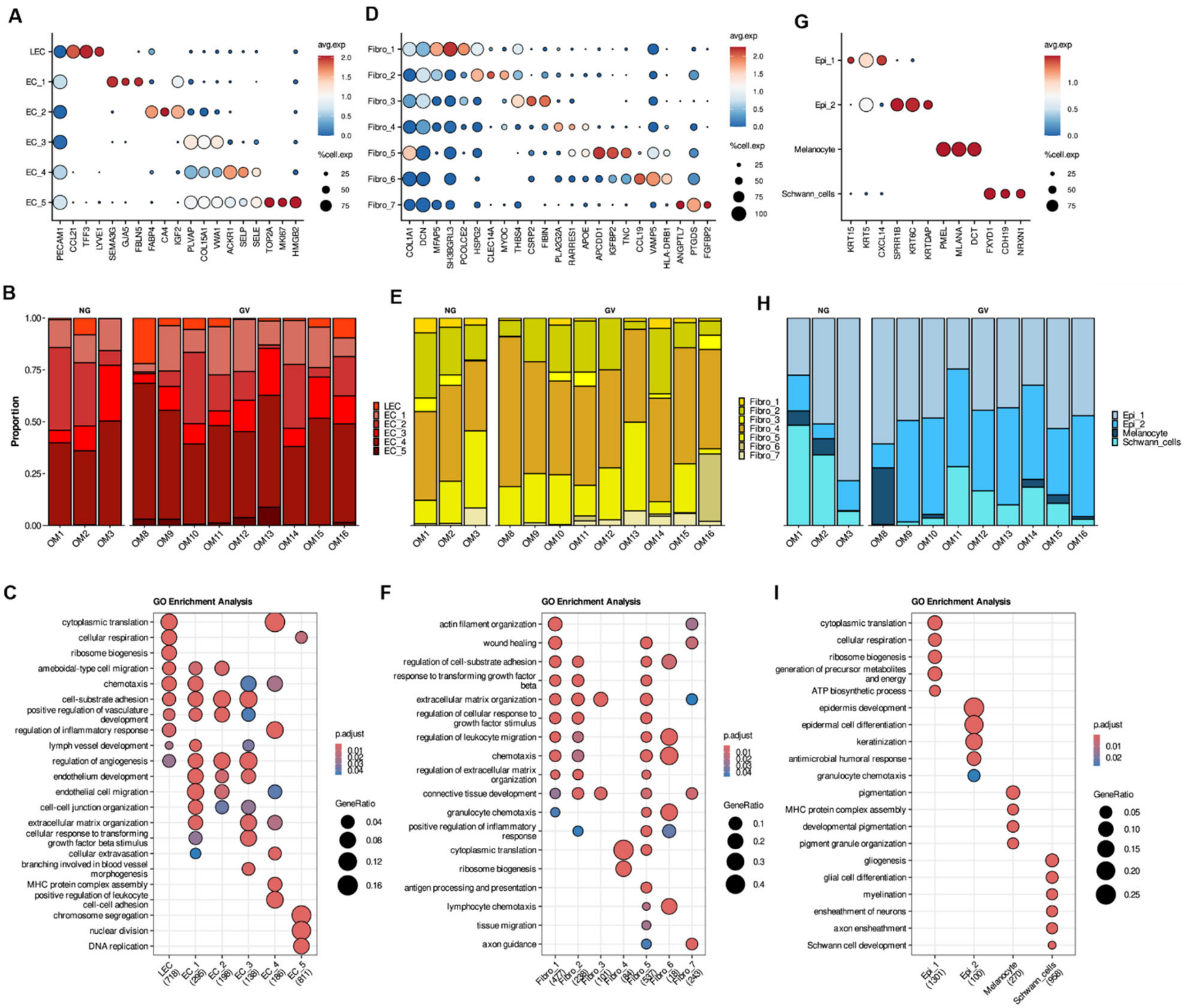

**Supplemental Figure 3. Expanded analysis of non-immune cells: Stromal and epithelial reprogramming in oral cGVHD. (A)** Dot plot of signature genes across endothelial subclusters, illustrating functional and phenotypic heterogeneity. **(B)** Proportional distribution of endothelial subclusters per individual patient. **(C)** GO Biological Process (GO BP) pathway enrichment analysis of pathways upregulated in endothelial subpopulations. Circle size indicates gene ratio; color scale reflects adjusted *p*-value (Benjamini–Hochberg). **(D)** Dot plot showing marker gene expression across fibroblast subclusters. **(E)** Cell proportions of fibroblast clusters per patient. **(F)** GO BP enrichment analysis for upregulated genes in fibroblast subpopulations. **(G)** Dot plot showing the expression of signature genes across epithelial clusters. **(H)** Bar plot of epithelial subcluster proportions per patient. **(I)** GO BP enrichment for epithelial subclusters. Size of each circle represents gene ratio, and color indicates adjusted $p$-value (one-sided Fisher's exact test with Benjamini–Hochberg correction).

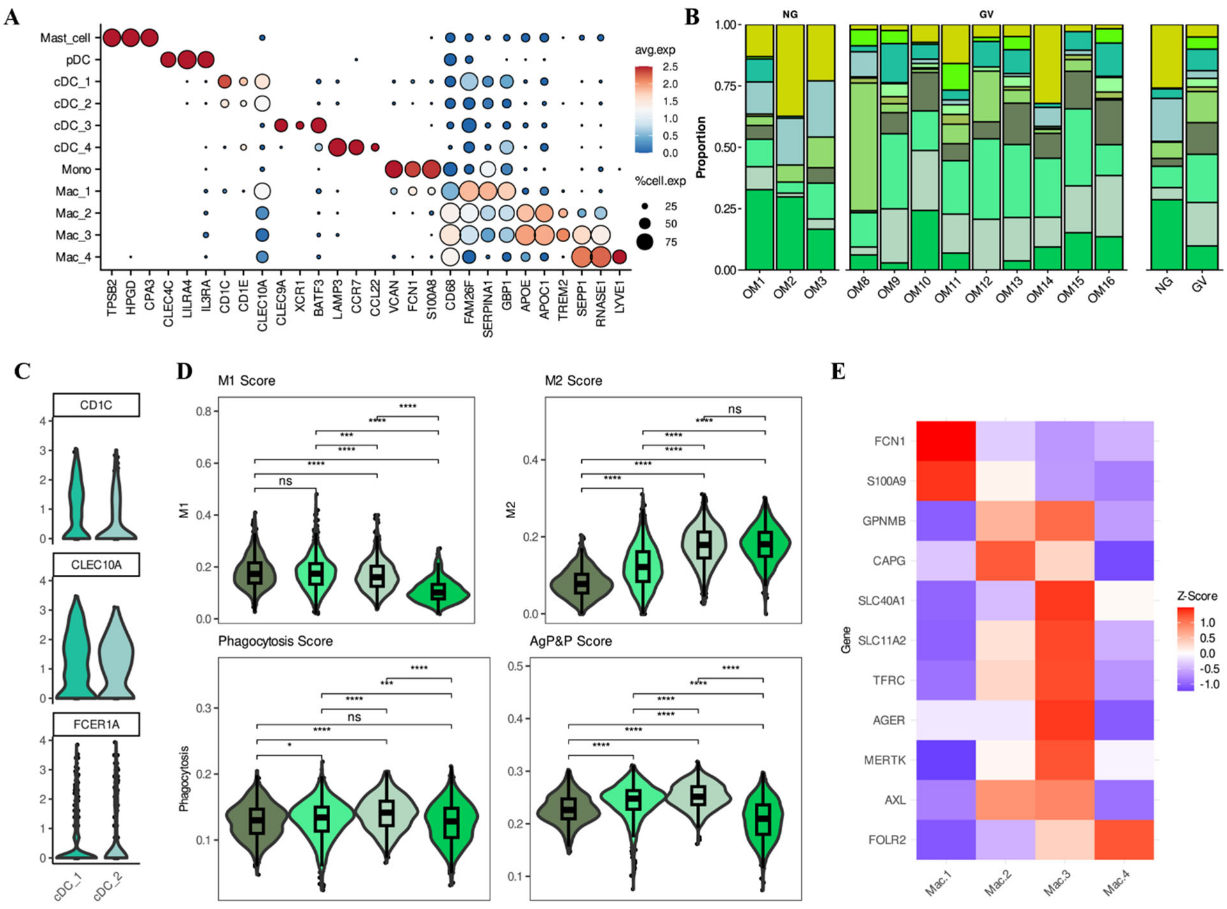

**Supplemental Figure 4. Extended characterization of myeloid subsets in oral cGVHD. (A)** Dot plot showing the expression of canonical marker genes used to annotate individual myeloid cell subsets, including dendritic cells, mast cells, monocytes, and macrophages. Dot size reflects the proportion of expressing cells; color intensity indicates average scaled expression. **(B)** Bar plots showing the proportion of each myeloid cluster across individual patients (left) and grouped by disease status (NG vs. GV, right). **(C)** Violin plots illustrating the expression of representative genes distinguishing cDC_1 and cDC_2 subsets. **(D)** Violin plots of M1, M2, antigen processing and presentation (AgP&P), and phagocytosis gene signature AUC scores across macrophage subsets. Statistical comparisons were performed using the Wilcoxon rank-sum test. **(E)** Heatmap showing expression patterns of selected marker genes across macrophage clusters, highlighting phenotypic heterogeneity.

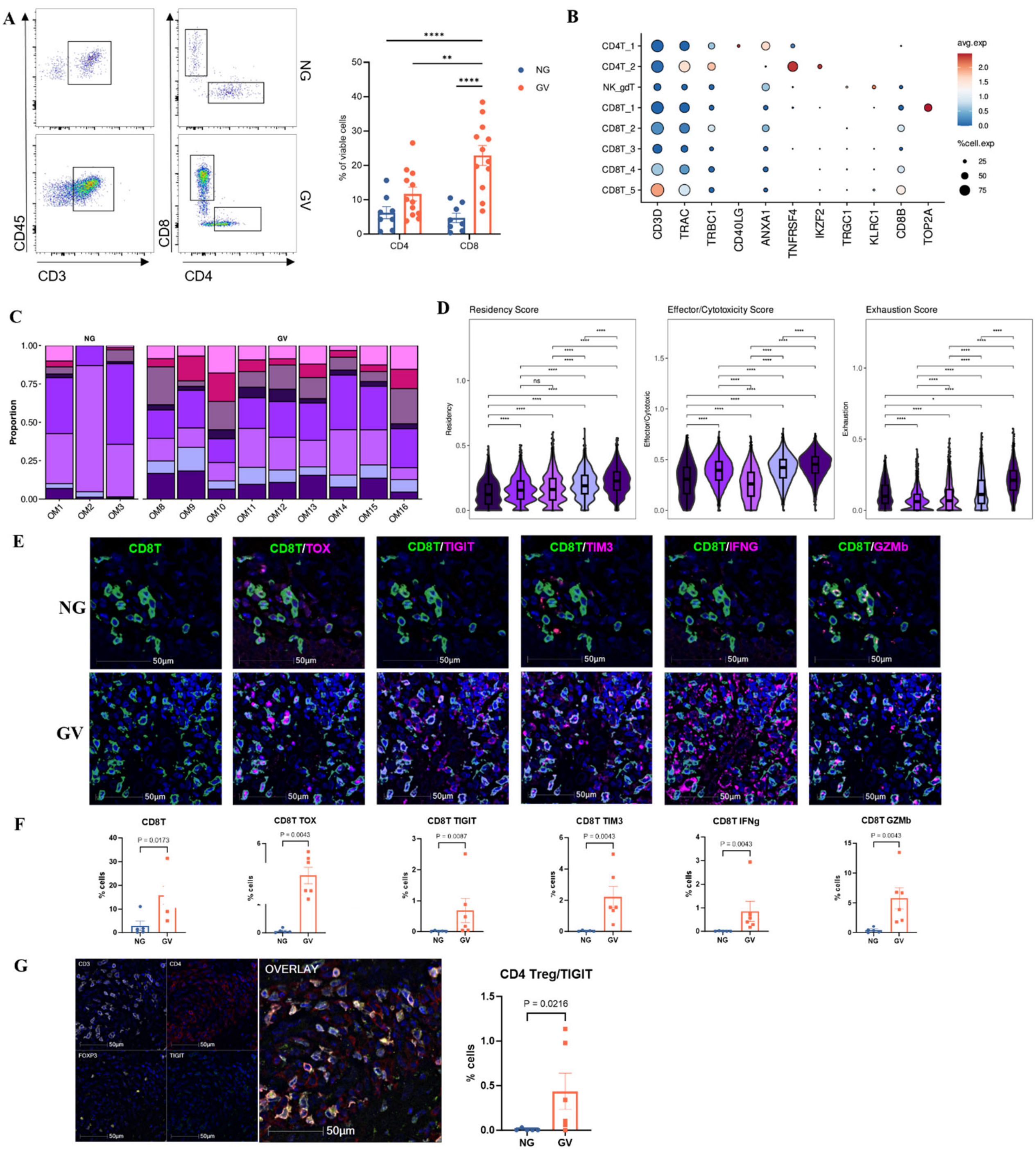

**Supplemental Figure 5. Additional characterization of exhausted-like CD8⁺ T cells and Tregs in oral cGVHD. (A)** Representative flow cytometry gating strategy for CD4⁺ and CD8⁺ T cell analysis from oral mucosa biopsies (left). Quantification of CD4⁺ and CD8⁺ T cell frequencies in NG (n = 8) and GV (n = 12) patients. Each dot represents an individual sample; bars indicate mean ± SEM. Statistical analysis performed using Two-way ANOVA, Sidak's post-test. **(B)** Dot plot showing expression of key signature genes across annotated lymphoid cell clusters. Dot size reflects the percentage of expressing cells; color intensity indicates scaled average expression. **(C)** Bar plots showing the relative proportions of lymphoid clusters across individual patients from NG and GV groups. **(D)** Violin plots display AUC scores for residency, cytotoxic/effector function, and exhaustion signatures across individual $CD8^+$ T cell clusters. Statistical significance assessed via Wilcoxon rank-sum test. **(E)** Representative IF images of OM tissue stained for CD8 (green), and exhaustion/cytotoxicity markers (TOX, TIGIT, TIM-3, IFNG, GZMB; magenta). DAPI (blue) marks nucleated cells. Scale bars = 50 μm. **(F)** Quantification of $CD8^+$ T cells and their co-expression with exhaustion/cytotoxicity markers in NG (n = 5) and GV (n = 6) tissues. Statistical significance determined by Wilcoxon rank-sum test. **(G)** Multiplex IF staining (left) of CD3 (white), CD4 (red), FOXP3 (yellow), and TIGIT (green) in oral mucosa tissue. DAPI (blue) indicates nuclei. Images acquired at 20× magnification; scale bars = 50 μm. Quantification (right) of $TIGIT^+$ $CD4^+FOXP3^+$ regulatory T cells (Tregs) in NG (n = 5) and GV (n = 6) samples. $p < 0.05$ considered statistically significant (Wilcoxon rank-sum test).

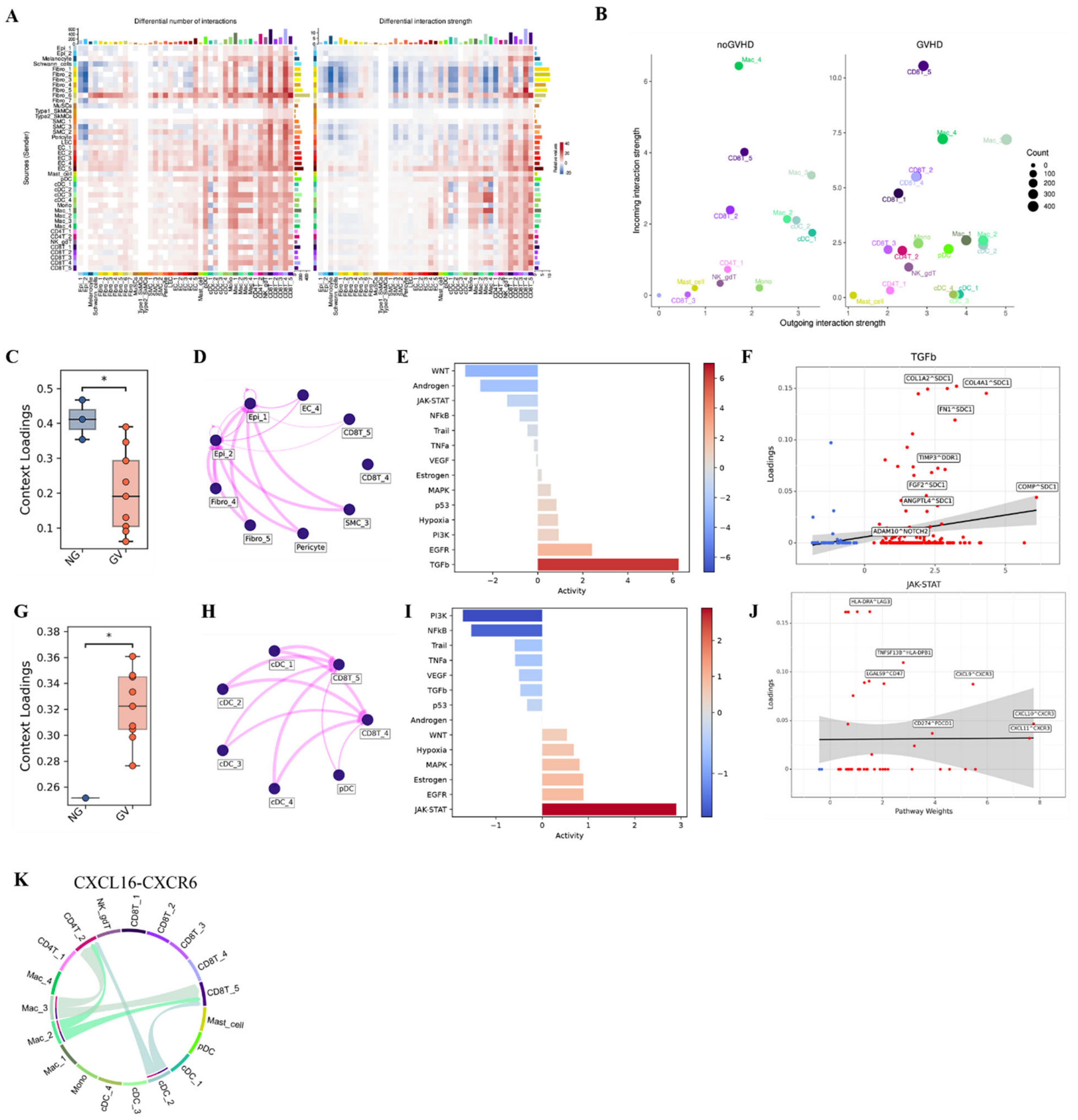

**Supplemental figure 6: Additional insights into stromal and immune cell signaling in the GVHD-affected oral mucosa microenvironment.** **(A)** Heatmaps showing the differential number (left) and strength (right) of predicted cell–cell interactions across cell types between NG and GV oral mucosa. **(B)** Scatter plots of outgoing versus incoming interaction strength per cell type, showing altered network topology in GV compared to NG patients. **(C)** Box plots compare downregulated context loadings between NG and GV patients related to stromal/epithelial-CD8$^{+}$

T cells interactions identified through Tensor-cell2cell analysis. Pairwise t-tests were performed, and p-values were adjusted using the Benjamini–Hochberg method. Statistical significance is indicated by asterisks ($p < 0.05$). **(D)** Network diagrams illustrate interactions between $CD8^+$ T cells and stromal/epithelial cells corresponding to (C), with connecting lines representing interaction strength. **(E)** Pathway enrichment analysis of ligand–receptor loadings identified using PROGENy. Color intensity reflects activity magnitude and reveals elevated TGF-β signaling activity in NG patients. **(F)** Scatter plots showing ligand–receptor pairs contributing to the TGF-β pathway enrichment shown in (E), with regression lines and confidence intervals (shaded areas) indicating trends and variability.

Table S1: Patient Demographics

| Demographics | scRNA seq | | |
|---|---|---|---|
| | NG | AG | GV |
| Number of cases | 3 | 4 | 9 |
| Age at sample, yr, median (range) | 25 (19-48) | 31 (21-40) | 29 (18-55) |
| Sex (F/M) | 1/2 | 0/4 | 5/4 |
| Race (%) | | | |
| White | 1 (33.3) | 1 (25) | 4 (44.5) |
| African | 2 (66.7) | 1 (25) | 1 (11.1) |
| Asian | | | 1 (11.1) |
| Hispanic | | | |
| Unknown | | 2 (50) | 3 (33.3) |
| Time post-HSCT, days, median (range) | 185 (182-186) | 179.5 (170-182) | 112 (64-550) |
| **Underlying Disease, n (%)** | | | |
| Lymphoma (HD/NHL) | 1 (33.3) | 2 (50) | 1 (11.1) |
| Acute leukemia (AML/ALL) | | 1 (25) | 4 (44.5) |
| Chronic leukemia (CLL/CML) | | | 1 (11.1) |
| MDS/myeloproliferative disorder (MDS, myelofibrosis, PV) | | | 2 |
| Sickle cell disease | 2 (66.7) | 1 (25) | |
| Other | | | 1 (11.1) |
| **Donor type, n (%)** | | | |
| Related | 2 (66.7) | 1 (25) | 8 (88.9) |
| Unrelated | 1 (33.3) | 3 (75) | 1 (11.1) |
| **HLA match, n (%)** | | | |
| Matched | | 1 (25) | 3 (33.3) |
| Mismatched | 1 (33.3) | 1 (25) | |
| Haploidentical | 2 (66.7) | 2 (50) | 6 (66.7) |
| **Sex match (donor -> recipient) , n (%)** | | | |
| F ® F | 1 (33.3) | | 3 (33.3) |
| F ® M | | | 2 (22.2) |
| M ® M | 2 (66.7) | 4 (100) | 2 (22.2) |
| M ® F | | | 2 (22.2) |
| Not Available | | | |
| **Conditioning regimen , n (%)** | | | |
| Myleoablative | 1 (33.3) | 1 (25) | 4 (44.4) |
| Nonmyeloablative | 2 (66.7) | 3 (75) | 5 (55.6) |
| History of TBI, n(%) | 1 (33.3) | 1 (25) | 3 (33.3) |
| **Stem cell source , n (%)** | | | |
| PBSC | 2 (66.7) | 3 (75) | 3 (33.3) |
| Bone Marrow | 1 (33.3) | 1 (25) | 6 (66.7) |
| History of aGVHD, yes, n (%) | 1 (33.3) | 2 (50) | 5 (55.6) |
| **Systemic immunosupressive treatment at time of saliva sample n (%)** | | | |
| None | | 2 (50) | 6 (66.7) |
| Steroid | | | 2 (22.2) |
| Calcineurin/mTOR inhibitor | 3 (100) | 1 (25) | 1 (11.1) |
| Calcineurin/mTOR inhibitor + Steroid | | 1 (25) | |
| Other | | | |

Table S2: 10X Chromium sequencing details

| S.No. | Disease group | Estimated No. of cells | Fraction Reads in cells | Mean Reads/cell | Median genes/cell | Total Genes detected | Median UMI/cell |
|---|---|---|---|---|---|---|---|
| 1 | NG | 2,782 | 62.50% | 84,354 | 1,862 | 22,633 | 5,326 |
| 2 | NG | 5,177 | 71.40% | 34,944 | 1,399 | 23,345 | 3,377 |
| 3 | NG | 1,955 | 76.70% | 51,601 | 1,944 | 22,011 | 4,449 |
| 1 | AG | 5,158 | 76.50% | 17,465 | 901 | 22,232 | 2,253 |
| 2 | AG | 4,096 | 56.70% | 55,262 | 1,217 | 23,127 | 2,930 |
| 3 | AG | 3,212 | 76% | 97,976 | 1,726 | 23,889 | 4,474 |
| 4 | AG | 992 | 60.30% | 143,743 | 955 | 20,466 | 4,153 |
| 1 | GV | 8,135 | 78.50% | 18,454 | 924 | 23,946 | 2,455 |
| 2 | GV | 2,266 | 54.60% | 54,649 | 1,184 | 22,427 | 2,982 |
| 3 | GV | 2,833 | 83.80% | 40,479 | 1,970 | 23,278 | 5,298 |
| 4 | GV | 2,499 | 76.20% | 60,154 | 1,876 | 22,852 | 5,138 |
| 5 | GV | 1,059 | 75.30% | 86,103 | 1,290 | 20,084 | 3,078 |
| 6 | GV | 10,379 | 80.90% | 25,996 | 1,299 | 24,597 | 2,817 |
| 7 | GV | 3,058 | 59.40% | 97,408 | 1,505 | 22,951 | 3,659 |
| 8 | GV | 2,269 | 59.70% | 71,716 | 1,447 | 21,541 | 3,649 |
| 9 | GV | 10,995 | 75.60% | 12,736 | 924 | 23,855 | 1,993 |

Table S3: Genes used to define cell state scores – non-immune cells (related to Figures 2D-F)

| Antigen Presentation Score | Chemotaxis score | Regulation of Inflammatory Response |
|---|---|---|
| GOBP_ANTIGEN_PROCESSING_AND_PRESENTATION (GO:0019882) | GOBP_LEUKOCYTE_CHEMOTAXIS (GO:0030595) | GOBP_REGULATION_OF_INFLAMMATORY_RESPONSE (GO:0050727) |

Table S4: Genes used to define cell state scores – myeloid cells (related to Figures 3H, I)

| M1 | M2 | Phagocytosis | AgP&P |
|---|---|---|---|
| IL1B | CCL17 | GOBP_PHAGOCYTOSIS (GO:0050766) | GOBP_ANTIGEN_PROCESSING_AND_PRESENTATION (GO:0019882) |
| IFNG | CCL22 | | |
| CD40 | CCL24 | | |
| CD80 | IL10 | | |
| CD86 | TGFB1 | | |
| IL1R1 | CCL13 | | |
| HLA-DRA | VEGFA | | |
| TLR2 | PDGFA | | |
| TLR4 | PDGFB | | |
| IFNAR2 | MMP9 | | |
| IFNAR1 | VEGFB | | |
| FCGR1A | TGFB2 | | |
| CCL3 | TGFB3 | | |
| CCL4 | MMP14 | | |
| CCL5 | MMP19 | | |
| CXCL8 | SEPP1 | | |
| CXCL9 | IDO2 | | |
| CXCL10 | CD163 | | |
| CXCL11 | MSR1 | | |
| TNF | LYVE1 | | |
| IL6 | STAB1 | | |
| IL1A | MARCO | | |
| KYNU | CD36 | | |
| IL12B | FCGR2A | | |
| SLC2A1 | IL1R2 | | |
| TFEB | IL4R | | |
| STAT1 | CD274 | | |
| NFKB1 | PDCD1LG2 | | |
| STAT3 | PDCD1 | | |
| IFITM3 | SIRPA | | |
| IRF3 | SIGLEC10 | | |
| IRF5 | LILRB1 | | |
| IRF7 | LILRB2 | | |
| SOCS3 | TEK | | |
| JUN | TREM1 | | |
| IDO1 | TREM2 | | |
| IRF1 | IL1RN | | |
| | STAT6 | | |
| | MAFB | | |
| | IRF4 | | |
| | SOCS1 | | |

Table S5: Genes used to define cell state scores – lymphoid cells (related to Figures 4D - F)

| Residency | Effector/Cytotoxicity | Exhaustion |
|---|---|---|
| CD69 | IFNG | HAVCR2 |
| ITGAE | GZMA | PDCD1 |
| ITGA1 | GZMB | CTLA4 |
| CXCR6 | GZMH | LAG3 |
| DUSP6 | PRF1 | TIGIT |
| RGS1 | CTSW | ENTPD1 |
| CRTAM | CST7 | CD244 |
| PRDM1 | NKG7 | BTLA |
| RUNX3 | GNLY | LAYN |
| ZNF683 | KLRK1 | CD160 |
| | KLRB1 | BATF |
| | KLRD1 | TOX |
| | HLA-DRB1 | |
| | HLA-DPA1 | |
| | EOMES | |
| | TNFSF9 | |

### Table S6: List of antibodies and reagents used in this study

| Antibodies | Description | Company | Catalog No. | Working oncentration | Experiment used | RRID number |
|---|---|---|---|---|---|---|
| CD45 | Rabbit polyclonal anti-CD45 | abcam | ab10558 | 1:100 | IF | AB_442810 |
| CCL4 | Rabbit polyclonal anti-CCL4 | Invitrogen | PA5-114961 | 1:200 | IF | AB_2899597 |
| CXCL13 | Goat polyclonal anti-CXCL13 | R&D | AF801 | 1:200 | IF | AB_355613 |
| CD8 | Mouse IgG2b anti-CD8 | BioRad | MCA1817T | 1:100 | IF | AB_323534 |
| CD3 | CoraLite® Plus 488-conjugated Mouse IgG2b anti-CD3 | ProteinTech | CL488-60181 | 1:100 | IF | AB_2883113 |
| IgG1 kappa Isotype Control (P3.6.2.8.1) | Alexa Fluor™ 488-conjugated Mouse IgG1,kappa | eBioscience | 53-4714-80 | 1:100 | IF | AB_470230 |
| Anti-Mouse IgG (H+L) | Cy™3 AffiniPure® Donkey Anti-Mouse polyclonal | Jackson Immunoresearch | 715-165-150 | 1:100 | IF | AB_2340813 |
| Anti-Rabbit IgG (H+L) | Alexa Fluor® 594 AffiniPure® Donkey Anti-Rabbit polyclonal | Jackson Immunoresearch | 711-585-1521 | 1:100 | IF | AB_2340621 |
| Anti-Goat IgG (H+L) | Alexa Fluor® 647 AffiniPure® Donkey Anti-Goat polyclonal | Jackson Immunoresearch | 705-605-003 | 1:100 | IF | AB_2340436 |
| DAPI | | ThermoScientific | 62248 | 1:2000 | IF | NA |
| CD8 | Anti-Hu CD8(C8/144B)-BX026 - Atto 550 | Akoya Biosciences | 4250012 | 1:100 | CODEX | AB_2915960 |
| TOX | Anti-Hu TOX(AKYP0098)-BX060 - Atto 550 | Akoya Biosciences | 240126 | 1:100 | CODEX | NA |
| LAG3 | Anti-Hu LAG3(AKYP0089)-BX055 - Alexa Fluor™ 647 | Akoya Biosciences | 4550058 | 1:100 | CODEX | AB_3096409 |
| IFNG | Anti-Hu IFNG(AKYP0093)-BX020 - Atto 550 | Akoya Biosciences | 4250062 | 1:100 | CODEX | AB_3476455 |
| CD4 | Anti-Hu CD4(AKYP0048)-BX003 - Alexa Fluor™ 647 | Akoya Biosciences | 4550112 | 1:100 | CODEX | AB_3094499 |
| FOXP3 | ANTI-HU FOXP3(AKYP0102)-BX031 - Alexa Fluor™ 647 | Akoya Biosciences | 4550071 | 1:100 | CODEX | AB_2927679 |
| TIGIT | Mouse IgG2a-BX002 - Atto 550 | BioLegend | A15153G | 1:100 | CODEX | NA |
| TIM3 | Goat polyclonal-BX030 - Cy5 | R&D | AF2365 | 1:50 | CODEX | AB_355235 |
| CD45 | Brilliant Violet 785™ anti-human CD45 | Biolegend | 304048 | 1:100 | Flow cytometry | AB_2563129 |
| CD3 | Alexa Fluor® 700 Mouse Anti-Human CD3 | BD Bioscience | 557943 | 1:200 | Flow cytometry | AB_396952 |
| CD4 | PerCP-Cyanine5.5 mouse anti-human CD4 (OKT4) | eBioscience | 45-0048-42 | 1:100 | Flow cytometry | AB_10804390 |
| CD8 | BV510 Mouse Anti-Human CD8 | BD Bioscience | 563256 | 1:200 | Flow cytometry | AB_2738101 |

| Reagents | | Company | Catalog No. |
|---|---|---|---|
| Collagenase P | | Millipore Sigma | 11213857001 |
| Deoxyribonuclease I from bovine pancreas | | Sigma- Aldrich | DN25-1G |
| gentleMACS™ C Tubes | | Miltenyi Biotech | 130-093-237 |
| Falcon™ Cell Strainers 40 µm | | Fischer scientific | 08-771-1 |
| RPMI-1640 | | Thermo Scientific | 61870127 |
| Fetal Bovine Serum | | ThermoFischer Scientific | A5256801 |
| Trypan Blue | | Invitrogen | T10282 |
| ACK Lysing buffer | | KD Medical | RGC-3015 |
| Xylene substitute | | Sigma- Aldrich | A5597-1GAL |
| Tris Base | | Sigma- Aldrich | T1378-500G |
| EDTA | | KD Medical | RGF-3130 |
| Tween 20 | | Affymetrix | 9005-64-5 |
| Fluoro-Gel (with Tris Buffer) | | Electron Microscopy Sciences | 17985-10 |
| UltraComp eBeads | | ThermoFischer Scientific | 01-2222-42 |
| Zombie UV Fixable Viability Kit | | Invitrogen | 423108 |
| Human TruStain FcX™ | | Biolegend | 422302 |
| Brilliant Stain Buffer | | BD Biosciences | 566349 |
| Paraformaldehyde 16% Aqueous Solution EM Grade | | Electron Microscopy Sciences | 15710 |
| Poly L-Lysine | | Sigma- Aldrich | P8920-100ML |
| CODEX Conjugation Kit | | Akoya Biosciences | 7000009 |
| CODEX Staining Kit | | Akoya Biosciences | 7000008 |
| 10X CODEX Buffer | | Akoya Biosciences | 7000001 |
| CODEX Assay Reagent | | Akoya Biosciences | 7000002 |
| Storage Buffer, 120mL | | Akoya Biosciences | 232107 |
| 96 well plates | | Akoya Biosciences | 7000006 |
| CODEX Nuclear Stain | | Akoya Biosciences | 7000003 |
| Codex Gaskets v2, pack of 10 | | Akoya Biosciences | 7000010 |
| 96 well plate seals | | Akoya Biosciences | 7000007 |
| coverslips | | Akoya Biosciences | 7000005 |
| BX030 - Cy5-RX030 | | Akoya Biosciences | 5350005 |
| BX002 - Atto 550-RX002 | | Akoya Biosciences | 5450023 |
| Histochoice clearing agent | | VWR | H103-4L |
| Ethanol, 100% solution (200 Proof) single bottle | | VWR | 95041-462 |
